# On the complexity of resting state spiking activity in monkey motor cortex

**DOI:** 10.1101/2020.05.28.121095

**Authors:** Paulina Anna Dąbrowska, Nicole Voges, Michael von Papen, Junji Ito, David Dahmen, Alexa Riehle, Thomas Brochier, Sonja Grün

**Affiliations:** Institute of Neuroscience and Medicine (INM-6 and INM-10) and Institute for Advanced Simulation (IAS-6), Jülich Research Centre, Jülich, Germany; RWTH Aachen University, Aachen, Germany; Institut de Neurosciences de la Timone, CNRS-AMU, Marseille, France; Theoretical Systems Neurobiology, RWTH Aachen University, Aachen, Germany

## Abstract

Resting state has been established as a classical paradigm of brain activity studies, mostly based on large scale measurements such as fMRI or M/EEG. This term typically refers to a behavioral state characterized by the absence of any task or stimuli. The corresponding neuronal activity is often called idle or ongoing. Numerous modeling studies on spiking neural networks claim to mimic such idle states, but compare their results to task- or stimulus-driven experiments, which might lead to misleading conclusions. To provide a proper basis for comparing physiological and simulated network dynamics, we characterize simultaneously recorded single neurons’ spiking activity in monkey motor cortex and show the differences from spontaneous and task-induced movement conditions. The resting state shows a higher dimensionality, reduced firing rates and less balance between population level excitation and inhibition than behavior-related states. Additionally, our results stress the importance of distinguishing between rest with eyes open and closed.

## Introduction

The resting state in behavioral studies is defined operationally as an experimental condition without imposed stimuli or other behaviorally salient events (***Raichle, 2009***; ***Snyder and Raichle, 2012***). It has become a classical paradigm for experiments involving large scale measurements of brain activity like fMRI and M/EEG (***Vincent et al., 2007***; ***Raichle, 2009***; ***Deco et al., 2011***; ***Snyder and Raichle, 2012***; ***Baker et al., 2014***). A major conclusion of these studies is that spontaneous brain activity in human and monkey can be characterized as a sequence of re-occuring spatio-temporal patterns of activation or deactivation resembling task-evoked activity, but present during rest (***Vincent et al., 2007***; ***Fox and Raichle, 2007***; ***van den Heuvel and Hulshoff Pol, 2010***) Defined on the *whole-brain level*, they are shaped by anatomical connectivity and derived from functional connectivity (***Honey et al., 2009***; ***Bastos and Schoffelen, 2016***).

While the exact link between the fMRI signal and neuronal activity is a matter of ongoing research (***Logothetis and Wandell, 2004***; ***Ekstrom, 2010***), resting state studies have also been carried out on the *level of single brain areas*. Here, the spontaneous activity is often referred to as ongoing, intrinsic, baseline, or resting state activity, and can be studied by means of, for example, optical imaging combined with single electrode recordings (***Arieli et al., 1996***; ***Tsodyks et al., 1999***; ***Kenet et al., 2003***). Such data, collected under anesthesia, were used to investigate the variability in evoked cortical responses (***Arieli et al., 1996***), the switching of cortical states (***Tsodyks et al., 1999***), and the link of these cortical states to the underlying functional architecture (***Kenet et al., 2003***). In our study, we aim to characterize the resting state on yet another spatio-temporal scale, namely on the *scale of simultaneous single unit (SU) spiking activity* recorded in macaque monkey (pre-)motor cortex. Spiking activity in monkey motor cortex has been studied extensively during arm movements, which gives rise to an increased average neuronal firing compared to wait (***Nawrot et al., 2008***; ***Rickert et al., 2009***; ***Riehle et al., 2018***). On a single unit level, direction-specific neuronal sub-populations encode the movement direction by firing rate modulations (***Georgopoulos et al., 1986***; ***Rickert et al., 2009***). These and other studies also investigated the spike time irregularity and the spike count variability in monkey motor cortex during various behavioral epochs: Movements have been related to a lower spike count variability across trials (***Rickert et al., 2009***; ***Churchland et al., 2010***; ***Riehle et al., 2018***) and to a higher spike time irregularity (***Davies et al., 2006***; ***Riehle et al., 2018***) compared to wait or preparatory behavior without movements. However, the resting state we analyze in this study is conceptually distinct from waiting or preparatory epochs: there is no task to prepare for and no signal to be anticipated. It is a state without any particular expectations or dispositions.

Studies on spiking neural network models aim to mimic brain dynamics down to the level of individual neuron activities. Bottom-up modeling approaches thereby derive the structure and parameters of network models from anatomy and electrophysiology, trying to reproduce and understand experimentally observed activity features of increasing complexity. For simplicity and due to missing knowledge of inputs to local circuits, such studies often focus on the idle state and intrinsically generated dynamics of networks (***van Vreeswijk and Sompolinsky, 1996, 1998***; ***Brunel, 2000***; ***Kumar et al., 2008***; ***Voges and Perrinet, 2010***; ***Potjans and Diesmann, 2014***; ***Dahmen et al., 2019***). This approach provides the prerequisite to study network dynamics and function in the presence of external stimuli and finally its relation to behavior. There are computational frameworks that can be used for a quantitative comparison or, ultimately, validation of such spiking neural network models against experiments (***Gutzen et al., 2018***), but data on single unit activity in resting-state condition is still lacking. As a consequence, network models are often compared to data collected in behavioral experiments, where tasks or stimuli lead to transient deviations from the resting-state statistics such as e.g. the average firing rates (***Georgopoulos et al., 1986***; ***Riehle et al., 1997***; ***Kaufman et al., 2013***; ***Riehle et al., 2018***). Recently developed biophysical forward modeling schemes (***Einevoll et al., 2013***) in combination with large-scale network models (***Potjans and Diesmann, 2014***; ***Schmidt et al., 2018a***,b; ***Markram et al., 2015***) in principle provide a possible way to employ existing resting-state fMRI and M/EEG data to benchmark spiking network models. However, the complexity of the dynamics on the level of individual neurons is lost in these mesoscopic and macroscopic measures of activity. Therefore, to provide a suitable reference for the validation of spiking neural network models on the single neuron (microscopic) level, we here present an analysis of massively parallel spiking activity in macaque monkeys at rest.

The aim of this study is a detailed characterization of the spiking activity at rest compared to task-induced and spontaneous movements. To this end, we recorded the ongoing activity with a 4×4 mm^2^ 100 electrode Utah Array (Blackrock Microsystems, Salt Lake City, UT, USA) situated in the hand-movement area of macaque (pre-)motor cortex. We performed two types of experiments:

1. During resting state experiments (REST), we recorded the neuronal activity of two monkeys seated in a chair with no task or stimulation. The spontaneous behavior was then classified into periods of (sleepy) rest and movements.
2. Reach-to-grasp experiments (R2G) (***Riehle et al., 2013***; ***Torre et al., 2016***; ***Brochier et al., 2018***; ***Riehle et al., 2018***) provide well-defined periods of task-related movements and task-imposed waiting. The latter behavior is similar to rest but contains a mental preparation task.

We ask if a distinction between spontaneous (resting and non-resting) and task-evoked (reach-to-grasp) neuronal dynamics is expressed on the level of single unit (SU) and network spiking activity. More specifically, we also ask if certain features of the neuronal firing during pure resting periods allow for a differentiation from spontaneous and task-induced movements, preparatory periods, or sleepiness. Contrary to this expectation, the motor system may show invariants, i.e., statistical properties of the neuronal spiking that do not change with respect to different behavioral epochs. While such comparisons have been performed on the level of local field potential (LFP) recordings, e.g., the investigation of behavior-related frequency modulations (***Engel and Fries, 2010***; ***Kilavik et al., 2013***), to our knowledge this is the first study to perform such comparison on the level of spiking activity.

In the following, we first detail how we performed the segmentation of REST recordings according to behavior, and then explored the activity of single neurons in different behavioral states. To investigate if there are comparable neuronal activity states in task-related data, we performed similar analyses for the R2G data. Apart from the SU dynamics, we also focused on network properties of the neuronal activities: We evaluated pairwise covariances, dimensionality of rate activities, and excitatory-inhibitory balance in the different behavioral states of both REST and R2G. The comparison to the R2G data enabled us to identify systematic network state changes which are less pronounced in REST.

## Results

We aim to determine in what regards spiking activity during rest is distinct from other behavioral states like spontaneous movements, sleepiness, movement preparation, or task-induced grasping. To do so, we first describe the behavioral segmentation of the REST data based on videos of the monkeys during the experiment, resulting in a sequence of defined behavioral states in REST. The segmentation of R2G was chosen as in previous studies on these data (***Riehle et al., 2018***). Then, we show that the behavioral segmentation is meaningful in terms of neuronal activity on two different scales: On the mesoscopic scale, which incorporates the collective behavior of neurons, we show that the LFP spectra differ across states. On the microscopic scale, we show that SU firing is correlated to the monkeys’ behavior, and examine the relation between behavior, spiking activity dimensionality and excitatory-inhibitory balance.

### Behavioral segmentation

Based on video recordings, each REST session (two per monkey) was segmented according to the monkey’s behavior, (cf. Materials and Methods: Behavioral Segmentation). Three states were considered: resting state (RS)—no movements and eyes open; sleepy resting state (RSS)—no movements and eyes (half-)closed; spontaneous movements (M)—movements of the whole body and/or limbs (Fig. 1A). For R2G recordings, two behavioral states were defined with respect to trial events. For these states, interval lengths of 500 ms were used: the first part of the preparatory period (PP)—500 ms after the first cue, when the monkey waits immobile for the GO; and a task-related movement period (TM)—an interval containing movement onset and grasping (Fig. 1B).

**Figure 1.**
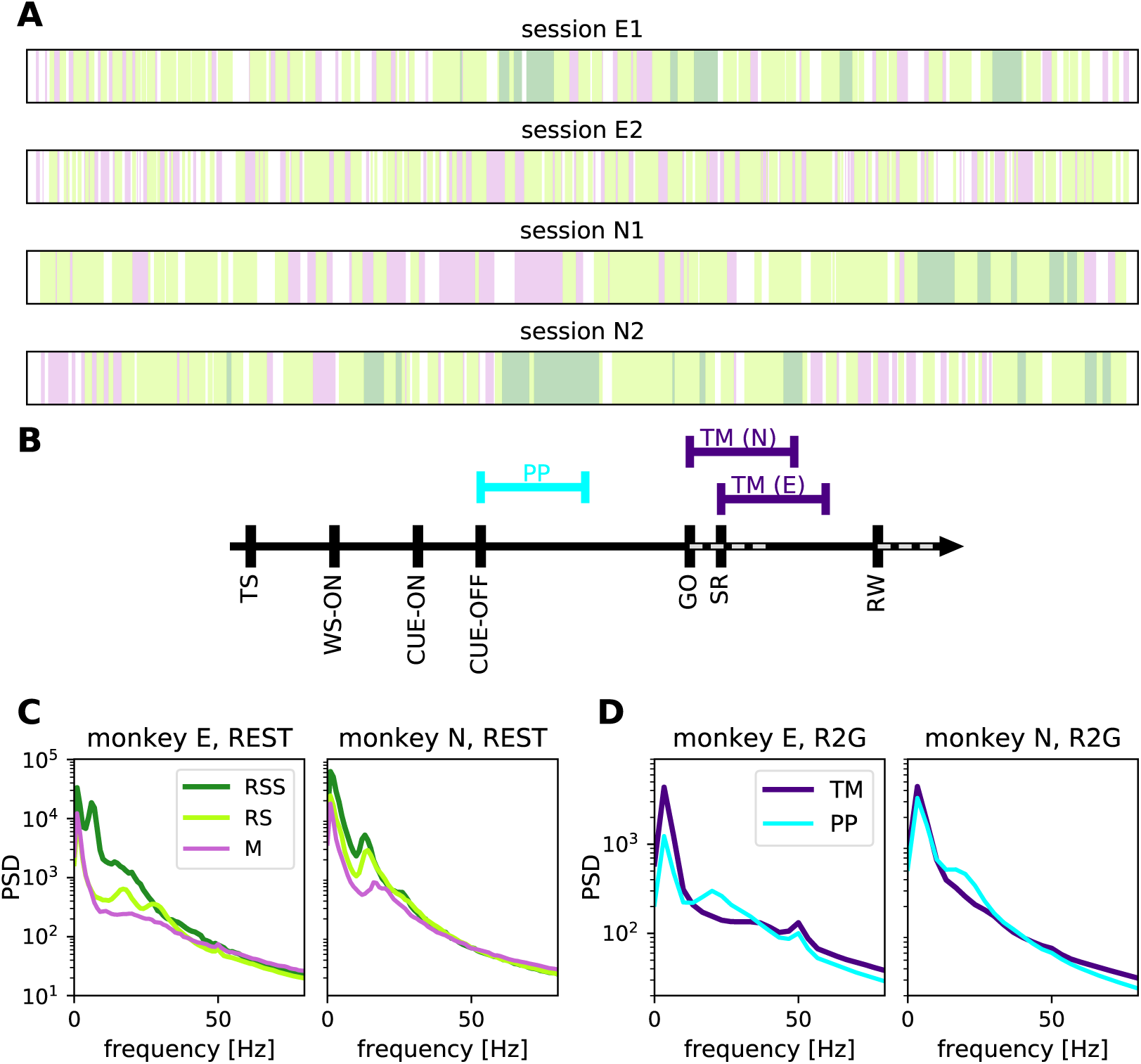
Behavioral segmentation in REST and R2G recordings. **(A)** Order and relative duration of the behavioral states defined within each REST session (single second precision): resting state (RS, light green) represents no movements and eyes open, sleepy rest (RSS, dark green) represents no movements and eyes (half-)closed, spontaneous movements (M, pink) represent movements of the whole body and limbs. **(B)** Order and timing of events within a single trial of an R2G session. Colored lines above the time axis indicate time intervals considered for the analysis: preparatory period (PP, cyan) and task-induced movements (TM, purple). SR indicates the switch-release event—beginning of the hand movement. PP was defined as [CUE-OFF, CUE-OFF+500 ms], and TM as [SR, SR+500 ms] for monkey E, and [SR-150 ms, SR+350 ms] for monkey N (different for the two monkeys due to differences in performance speed). **(C and D)** Power spectral density of LFP in different behavioral states. Panels in C pertain to REST, panels in D to R2G, left for monkey E and right for monkey N, respectively. States are defined in A and B. The peak at 50 Hz in the R2G spectra is an artifact (line frequency) and was not considered.

The visual segmentation is substantiated by comparison of the LFP spectra in the above defined states (Fig. 1C). The relationship between LFP and behavior has been shown in several studies, e.g. ***Pfurtscheller and Aranibar (1979)***; ***Fontanini and Katz (2008)***; ***Engel and Fries (2010)***; ***Takahashi et al. (2011)***; ***Kilavik et al. (2013)***. Beta oscillations (≈13 to 30 Hz) have been linked to states of general arousal, movement preparation, or postural maintenance (***Baker et al., 1999***; ***Kilavik et al., 2012***) and are typically suppressed during active movement (***Pfurtscheller and Aranibar, 1979***).

In our data, RS and PP show peaks in the range from ≈10 to ≈30 Hz (alpha/beta range), the peak in PP occurs for a higher frequency than in RS. In both monkeys, M and TM contain more power compared to other states in frequencies above ≈50 Hz (gamma), while beta power is reduced. However, the spectrum during RSS differs between monkeys. In monkey E, RSS seems to be a distinct physiological state: it shows strong slow oscillations, as to be expected (***Gervasoni et al., 2004***; ***Fontanini and Katz, 2008***) for a sleepy version of RS. In monkey N, however, the spectra during RSS are more similar to RS, but still with more power in the lower frequency bands.

Table 1 lists all single recording sessions for both REST and R2G experiments. It provides information about the number of SUs (separated into putative excitatory / broad spiking (bs) and putative inhibitory / narrow spiking (ns)) and the number of data slices (3 or 0.5 s long in REST) or trials (R2G), in the different behavioral states. Thus in total we have 2 sessions of 15-20 min from each monkey during REST, and 6/5 sessions (of similar durations) of monkey E/N during R2G. This results in 627 R2G trials of monkey E and 635 trials of monkey N. These were compared to the following numbers of data segments of 0.5 s during REST: 232, 2028 and 508 segments of RSS, RS and M, respectively, for monkey E and 372, 1676 and 492 segments for monkey N. For details on cutting the data see Materials and Methods: Behavioral Segmentation.

**Table 1.** List of all considered experimental recordings. Session names (first column) starting with “e” refer to monkey E, and with “i” to monkey N. Throughout the manuscript the REST sessions are referred to as E1, E2, N1 and N2. Each R2G trial yields one PP and one TM period, equally long (0.5 s each).

| REST session | #slices: 3 s (0.5 s) |  |  | #SUs (#bs, #ns) |
| --- | --- | --- | --- | --- |
|  | RSS | RS | M |  |
| e170103-002 (E1) | 40 (232) | 196 (1058) | 43 (200) | 115 (56, 50) |
| e170131-002 (E2) | 0 | 189 (970) | 67 (308) | 133 (67, 56) |
| i140701-004 (N1) | 20 (114) | 151 (840) | 58 (312) | 130 (76, 45) |
| i140615-002 (N2) | 45 (258) | 156 (836) | 36 (180) | 154 (78, 62) |
| R2G session | #trials |  |  | #SUs (#bs, #ns) |
| e161212-002 | 108 |  |  | 131 ( 50, 56) |
| e161215-001 | 102 |  |  | 92 ( 41, 37) |
| e161220-001 | 114 |  |  | 98 ( 45, 33) |
| e161222-002 | 102 |  |  | 118 ( 57, 41) |
| e170105-002 | 101 |  |  | 115 ( 60, 45) |
| e170106-001 | 100 |  |  | 116 ( 57, 43) |
| i140613-001 | 93 |  |  | 138 ( 71, 56) |
| i140616-001 | 130 |  |  | 154 ( 75, 61) |
| i140617-001 | 129 |  |  | 154 ( 83, 59) |
| i140703-001 | 142 |  |  | 142 ( 84, 46) |
| i140704-001 | 141 |  |  | 124 ( 70, 43) |

### Relation between neuronal firing and behavior

A prerequisite for the following analyses is to formalize a relationship between neuronal spiking activity and the behavioral states of a monkey. Therefore, we quantified the correlation between SU firing and behavior. This is by no means to be taken as a decoding approach, but rather as a substantiation for the approach taken above to differentiate between behavioral states in REST.

Figure 2A shows the time-resolved firing rates (FR) of all recorded SUs in one REST session (N1) (Sec. Materials and Methods: Behavioral correlation). They change in time and are variable across SUs, which is true for all REST sessions. The firing rates range from 0 up to ≈100 spikes per second. Some SUs exhibit a consistent firing (not visible by eye), e.g., unit 4 in Fig. 2A with a small absolute standard deviation, FR= 1.23 ± 1.16, and similarly unit 127 (relative standard deviation FR= 25.29± 6.46). The firing of other SUs changes considerably over time, e.g., unit 126 with a large absolute standard deviation FR= 20.08 ± 12.15, and similarly unit 17 with a relative standard deviation FR= 1.74± 3.53.

**Figure 2.**
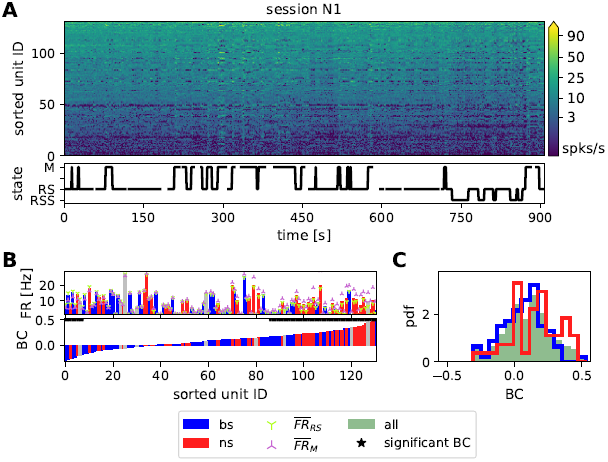
The correlation between SU firing and behavior for one REST session of monkey N. **(A)** Time- and population-resolved firing (spikes/s). SUs are sorted according to average firing rates in increasing order from bottom to top. The state vector describing the monkey’s behavior is shown below. The time resolution is 1 s. Empty spaces denote periods of unclassified behavior, vertical lines indicate transitions between identified states. **(B)** Comparison of average firing rates and behavioral correlation (only M and RS are taken into account). The SUs in both diagrams are sorted according to increasing values of BC. Red bars indicate broad-spiking (bs, putative excitatory) and blue narrow-spiking (ns, putative inhibitory) SUs, grey indicates unclassified units. Green and pink triangles on top of the FR bars indicate the average firing rate of the corresponding SU during RS and M, respectively. Black stars above the BC bars indicate significant correlations. **(C)** Distributions of BC values. In this recording session the difference between the ns (red) and the bs (blue) distribution is significant.

To examine this variability with respect to the behavior of the monkey, we defined a behavioral state vector (cf. bottom panel of Fig. 2A). Its entries represent the behavioral states: the value is set to +1 if there are movements (M) and −1 if the monkey is at rest (RS), and for the following analysis all other states are not taken into account. The bottom row of Fig. 2B shows the values of the behavioral correlation (BC, see Sec. Materials and Methods: Behavioral correlation) between the state vector and the firing rate (in 1 s bins) of each of the SUs, ordered from minimum to maximum. The panel above shows the FR of the corresponding SUs in identical order, averaged over the whole recording period (bars), only over RS periods (green markers) and only over M periods (pink markers). Most SUs increase their firing rate during M (mostly on the right side of the panel), many of them significantly (BC *>* 0.17, *p<* 0.001). A much smaller set of SUs increases the firing rate during RS (BC *<* –0.17), seen mostly on the left side of the panel. This asymmetry between the two states is reflected by the positive average BC in all 4 REST sessions (Tab. 2 col. 3). The second column of Tab. 2 lists the percentage of SUs with significant BC, for all sessions (see Sec. Materials and Methods: Behavioral correlation for the derivation of the BC significance). They range from 40.8 to 66.9%, however, neither the sign nor the amount of the behavioral correlation can be reliably predicted from the average FR: Both SUs with very high or very low mean FR show negative, positive and close to zero BC values. This is also indicated by the insignificant correlation between FR and BC (Tab. 2 col. 4): *ρ*_BC,FR_. Yet, the consistently negative *ρ*_BC,FR_ values suggest that SUs with smaller firing rates tend to be more sensitive to behaviour that highly active ones (the lower the mean FR, the higher the mean BC).

**Table 2.** Behavioral correlation for all REST sessions. The second column gives the percentage of SUs that show a significant behavioral correlation (BC, *p<* 0.001), the third column gives BC averages, and the fourth column the Spearman rank correlation between average SU firing rate and BC (*ρ*_BC,FR_). Column five lists the percentages of SUs that change their firing significantly (*p<* 0.001) with the behavioral state, obtained with a Kruskal-Wallis test on M, RS, and RSS. The numbers in brackets indicate the values obtained when separating between bs (first entry) and ns (second entry) SUs. 1 s resolution.

| Session | % SUs<sub>BC</sub> (bs, ns) | mean BC (bs, ns) | $\rho_{BC,FR}$ | % SUs<sub>KW</sub> (bs, ns) |
| --- | --- | --- | --- | --- |
| E1 | 53.9 (50, 60) | 0.13 (0.14, 0.13) | -0.19, $p=0.03$ | 54.7 (51.7, 58) |
| E2 | 66.9 (64.2, 71.4) | 0.14 (0.14, 0.15) | -0.09, $p=0.3$ | 76.9 (79.1, 76.8) |
| N1 | 40.8 (32.9, 53.3) | 0.10 (0.07, 0.15) | -0.13, $p=0.2$ | 48.1 (39, 62.2) |
| N2 | 46.8 (39.7, 54.8) | 0.12 (0.09, 0.14) | -0.04, $p=0.7$ | 43.5 (39.7, 50) |

In order to include the RSS state (in addition to M & RS) in the correlation of neuronal activity and behavioral states, we performed a Kruskal-Wallis test (KW) per SU, which provides information about the significance, but no quantification of the strength of the correlation. The obtained percentage of significantly correlated SUs (*p<* 0.001) ranges from 55% to 77% in monkey E and from 44% to 48% in monkey N (last column of Tab. 2). Thus, we find a clear inter-relation of the behavioral state and the neuronal activity.

Since firing rates seem to be not indicative of the behavioral state, we further differentiated the data into putative excitatory and inhibitory neurons (Sec. Materials and Methods: Pre-processing). Fig. 2C shows the distribution of BC values obtained in session N1 for all SUs (green shaded area), and for SU separated into putative excitatory / broad spiking SUs (bs) and putative inhibitory / narrow spiking (ns) SUs (blue and red lines, respectively). In this session we find a significant difference between the ns and the bs BC distribution. However, this could not be substantiated in the data from other recording sessions, indicating that the neuron type does not determine the strength of correlation with behavior. Still, firing of putative inhibitory as compared to excitatory neurons seems to be more related to behavioral states. This is indicated by higher percentages of significantly correlated ns than bs SUs (cf. Tab. 2 col. 2&5), particularly in monkey N, see also the higher mean BC of ns in monkey N (cf. Tab. 2 col. 3).

In order to include also the RSS state (in addition to M & RS) in the correlation of neuronal activity and behavioral states, we performed a Kruskal-Wallis test (KW) per SU, which provides information about the significance, but no quantification of the strength of the correlation. The obtained percentage of significantly correlated SUs (*p<* 0.001) ranges from 55% to 77% in monkey E and from 44% to 48% in monkey N (last column of Tab. 2). Thus, we find a clear inter-relation of the behavioral state and the neuronal activity of the observed population.

To examine in more detail behavior-related modulations of average FR, we performed a set of pairwise comparisons between behavioral states per SU (using 3 s slices, see Sec. Materials and Methods: Behavioral correlation). Table 3 summarizes the results by listing the percentages of SUs that significantly change their FR with respect to behavior. We observe that ≈34 to 67% of the SUs show significantly higher FR during M as compared to RS, but still, 5 to 11% of SUs show significantly higher FR during RS (second and fifth column in Tab.3). Correspondingly, the percentages for RSS versus M show a similar tendency (≈25 to 48% and 2 to 8%, respectively, col. 3&6). This confirms the results obtained so far, i.e., that there are mostly lower firing rates during rest (RS and RSS) than during movement (M).

**Table 3.** Pairwise comparisons of SU firing rates in different states. Percentage of SUs that exhibit significantly lower (first three columns) or higher (last three columns) firing rates in the first of the two states indicated in the column header (RS vs M, RSS vs M, and RS vs RSS) for all REST sessions (in 3 s slices).

| Session | RS < M | RSS < M | RS < RSS | RS > M | RSS > M | RS > RSS |
| --- | --- | --- | --- | --- | --- | --- |
| E1 | 37.6 | 24.8 | 18.8 | 7.7 | 7.7 | 3.8 |
| E2 | 67 |  |  | 11 |  |  |
| N1 | 33.6 | 30.5 | 3.8 | 6.1 | 1.5 | 19.8 |
| N2 | 42.9 | 48.1 | 2.6 | 4.5 | 1.9 | 30.5 |

The properties of M in relation to RS are consistent in the two monkeys, but the RSS state differs between them. Only 3 to 4% of SUs show significantly lower firing in RS than in RSS in monkey N, but in monkey E the percentage is ≈20% (Tab. 3 col. 4). Vice versa, only 3.8% of SUs show significantly higher firing in RS than in RSS in monkey E, while it is ≈20 to 30% in monkey N (last column). Moreover, the percentages of SUs showing lower firing during RS and RSS as compared to M (second and third column in Tab.3) are rather similar in monkey N. However, in monkey E only 25% of SUs show higher firing during M than during RSS while 38% of the SUs show a higher firing during M as compared to RS. Thus, in agreement with our observations of the LFP spectra (cf. Sec. Results: Behavioral segmentation), rest and sleepy rest in monkey E express rather different features while they are quite similar for monkey N.

Above we show that the firing of approximately half of the SUs is significantly correlated to the behavior and that RS is, on average, associated with lower FRs than movements. However, the absolute value of the FR alone is not predictive of the response of a SU to different behavioral states. In the following section, we aim to investigate other aspects than mere SU spiking in different behavioral states.

### Further single unit firing properties and their relation to behavior

Given the relation between behavior and SU firing rate modulations, we now ask if other features of SU activity can be directly linked to particular behavioral states. From now on, we include R2G data to additionally look for differences on the level of spontaneous versus task-related behaviors. The box plots in Fig. 3 and the values listed in Tab. 5 describe averaged firing rates (FR), local coefficients of variation (CV2) and the Fano factor (FF), calculated for 0.5 s time slices of all REST and R2G sessions, per SU and time slice (see Sec. Materials and Methods: Data analysis). The CV2 characterizes the (ir-)regularity of neuronal firing across time. A value closer to zero (CV2⪅0.5) indicates regular spiking, Poissonian firing is characterized by CV2=1 (***Shinomoto et al., 2003***; ***Voges and Perrinet, 2010***), and values higher than one indicate more irregular spiking. The FF describes the variability of SU spike counts across trials (R2G) or time slices (REST) (***Nawrot et al., 2008***; ***Nawrot, 2010***; ***Riehle et al., 2018***). It equals one for a Poisson process and decreases for more reliable spiking.

**Figure 3.**
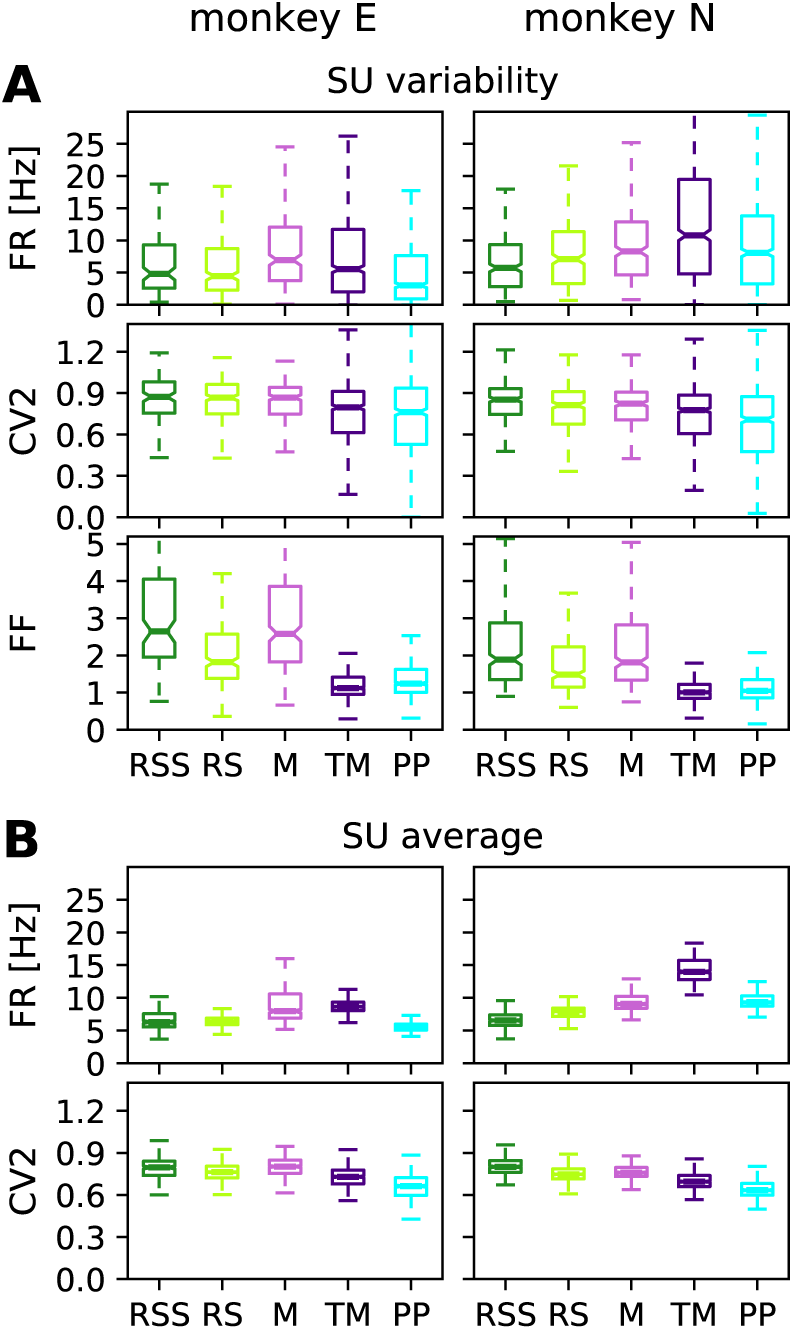
Comparison of firing properties in REST & R2G states calculated for 0.5 s data slices. **(A)** Box plots showing the variability across SUs: firing rate, spiking regularity, and spike count variability characterized by FR, CV2, and FF, here averaged over time slices. **(B)** Box plots showing the variability over time: distributions of time-resolved FR and CV2 averaged over SUs. Data pooled over REST sessions, two for each monkey (states RSS, RS, and M), and over R2G sessions, six of monkey E and five of monkey N (states TM and PP).

Averaged across time slices (Fig. 3A), FR shows the highest median in movement states (M & TM), while it is lower in RS(S) and PP (the differences being mostly significant, see below). Inferred from CV2, the firing is less regular in REST as compared to R2G states, and slightly less regular during TM than during PP, both showing a larger spread of values than the REST states. These differences are minor compared to the differences in the spike count variability: R2G states exhibit a much smaller and less variable FF, i.e., a higher reliability. M and RSS show the highest spread of FF, i.e. highest SU variability. The RS state exhibits a medium mean and spread of FF values.

Kruskal-Wallis tests on all 5 behavioral conditions yield highly significant differences between states for each measure (*p*≪ 0.0001) for both monkeys. The results of all pairwise comparisons are listed in Tab. 4. For both monkeys, most differences are significant, though CV2 differentiates primarily between REST and R2G recordings, thus between spontaneous and task-related behaviors.

**Table 4.** Significance of pairwise comparisons of firing rate (FR), local coefficient of variance (CV2) and Fano factor (FF) results shown in Fig. 3A. Upper triangle of each table: monkey E; lower: monkey N. Stars indicate significant differences (*: *p<* 0.001) and minuses insignificant differences after Bonferroni correction with *α* = 0.001. Grey background highlights significant results.

| FR | RSS | RS | M | TM | PP |
| --- | --- | --- | --- | --- | --- |
| RSS |  | - | - | - | * |
| RS | - |  | * | - | * |
| M | * | - |  | - | * |
| TM | * | * | * |  | * |
| PP | * | - | - | * |  |

| CV2 | RSS | RS | M | TM | PP |
| --- | --- | --- | --- | --- | --- |
| RSS |  | - | - | * | * |
| RS | - |  | - | * | * |
| M | - | - |  | * | * |
| TM | * | - | - |  | - |
| PP | * | * | * | * |  |

| FF | RSS | RS | M | TM | PP |
| --- | --- | --- | --- | --- | --- |
| RSS |  | * | - | * | * |
| RS | * |  | * | * | * |
| M | - | * |  | * | * |
| TM | * | * | * |  | * |
| PP | * | * | * | - |  |

**Table 5.** Quantification of average firing rate (FR, top row), regularity in spiking (CV2, middle row), and spike count variability (FF, bottom row). Given are mean values (averaged across time slices and SUs) and corresponding standard deviations with respect to SUs. All values are obtained from 0.5 s slices, for different behavioral states in REST (RSS, RS, M) and R2G (TM, PP), pooled across all recordings of the respective type.

| Monkey | RSS | RS | M | TM | PP |
| --- | --- | --- | --- | --- | --- |
| Firing rate FR [Hz] |  |  |  |  |  |
| E | $6.60 \pm 5.34$ | $6.43 \pm 5.74$ | $8.79 \pm 6.57$ | $8.74 \pm 9.69$ | $5.65 \pm 6.92$ |
| N | $6.64 \pm 4.66$ | $7.81 \pm 5.51$ | $9.44 \pm 6.32$ | $14.34 \pm 12.90$ | $9.58 \pm 7.74$ |
| Local coefficient of variation CV2 |  |  |  |  |  |
| E | $0.86 \pm 0.16$ | $0.83 \pm 0.17$ | $0.83 \pm 0.15$ | $0.78 \pm 0.31$ | $0.76 \pm 0.36$ |
| N | $0.83 \pm 0.15$ | $0.79 \pm 0.15$ | $0.80 \pm 0.16$ | $0.74 \pm 0.24$ | $0.69 \pm 0.29$ |
| Fano factor FF |  |  |  |  |  |
| E | $3.14 \pm 1.53$ | $2.06 \pm 0.88$ | $3.05 \pm 1.72$ | $1.32 \pm 0.82$ | $1.41 \pm 0.64$ |
| N | $2.31 \pm 1.30$ | $1.86 \pm 1.07$ | $2.29 \pm 1.42$ | $1.12 \pm 0.64$ | $1.21 \pm 0.78$ |

Averaging across SUs (Fig. 3B), we examine the variability in time. Note that even though the number of RS time slices highly exceeds that of SUs (cf. Tab. 1), the observed spread of the corresponding values is much smaller. This holds for all behavioral states. Since the variability across time slices (panel B) is much smaller than the variability across SUs (panel A), we later on averaged over time and considered only the variability with respect to SUs.

In summary, we find high SU variability in most of the measures for most of the states and the observed differences between states are mostly significant. Resting periods are characterized by rather low firing rates as compared to movements in agreement with the results in Sec. Results: Relation between neuronal firing and behavior. The RS in particular shows a higher reliability (lower FF) than M and RSS, but all REST states show a clearly higher FF as compared to R2G states.

### Network firing properties

We now turn towards the analysis of coordinated firing as opposed to single unit dynamics. Co-ordination between neurons can be measured at various time scales and quantified with various methods. We here consider spike-count covariances calculated for 3 s slices with a bin size of 100 ms, see Sec. Materials and Methods: Covariances and dimensionality. To this end, we first show the covariance (COV) distributions, averaged over slices of the REST data (Fig. 4A). While the average value during all REST behaviors is close to zero, the spread of the COV distributions differs between states, leading to highly significant differences (*p* ≪ 0.0001). In monkey E, the standard deviation of the covariances is considerably lower during 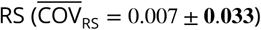 than during 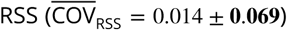 and 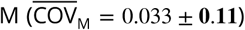. The same is true for monkey N: 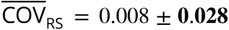 compared to 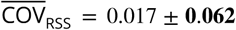 and 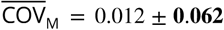. Statistical comparison of the shape of the distributions with two-sample Kolmogorov-Smirnov tests reveals significant differences for all pairs in both monkeys. In summary, we find that neuronal firing is less correlated during rest as compared to movements and we again observe distinct RSS properties in the two monkeys.

**Figure 4.**
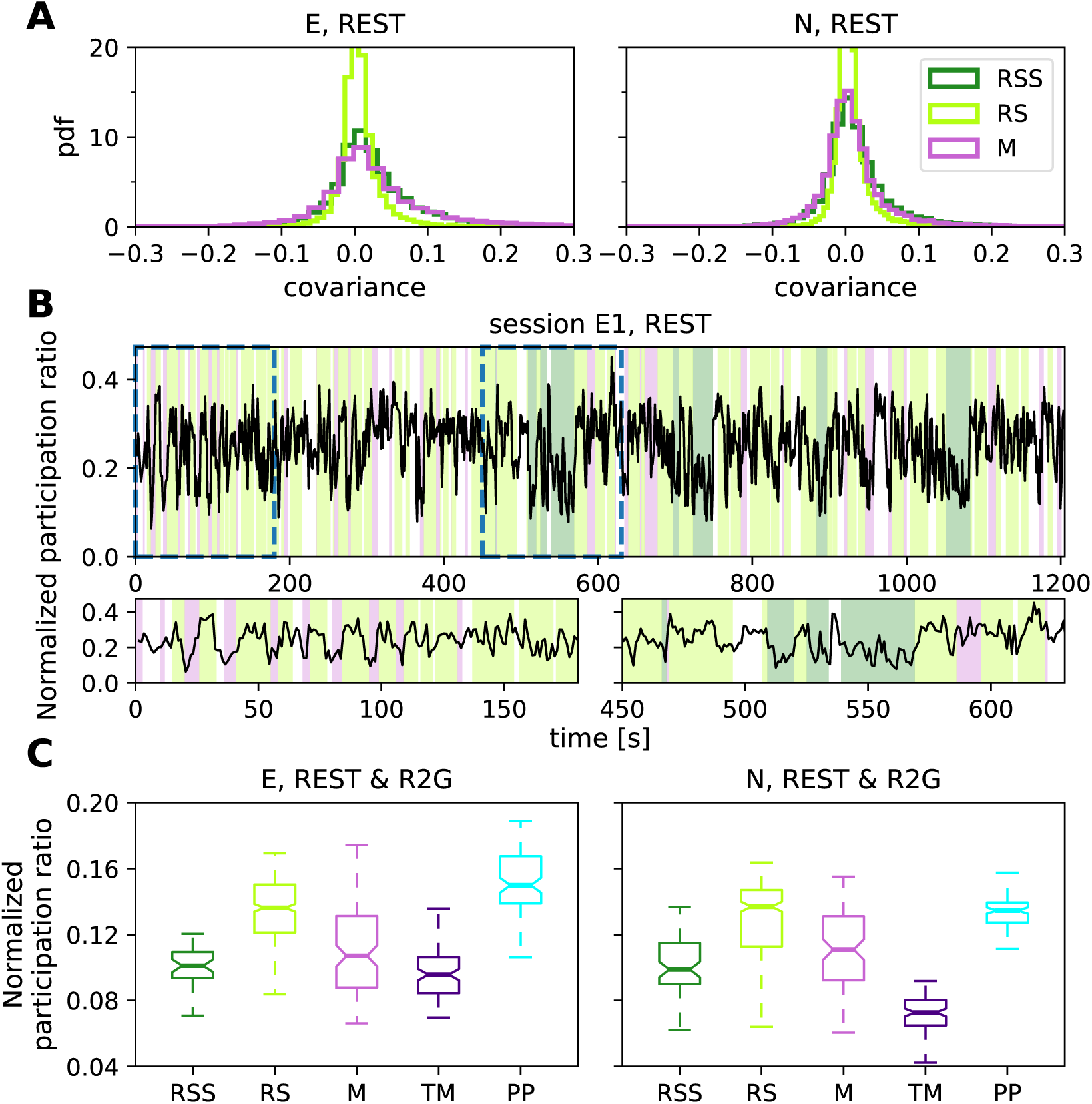
Network firing properties. **(A)** Distributions of pairwise covariances (COV) for the REST recordings of monkey E (left) and monkey N (right), calculated in 3 s slices with 100 ms bins, averaged over slices per SU pair and pooled over sessions. **(B)** Time-resolved participation ratio in session E1, calculated in 3 s long sliding windows with an overlap of 2 s. Each value on the plot corresponds to the center of the respective window. Colors in the background indicate behavioral states (cf. legend in the right panel of A). Two bottom panels show close-up view at periods marked by dashed lines in the top panel. **(C)** Dimensionality: Box plots show the normalized participation ratio of REST (RSS, RS, M) and R2G (TM, PP) states for monkey E (left) and monkey N (right), each single value of the distributions corresponds to a single 3 s data slice. Pooled over sessions.

The differences in the COV distributions motivate a more detailed investigation of the coordination of all recorded neurons. Apart from mean and variance of the covariance distribution, another summarizing measure for the covariance structure has been established and discussed in the recent years: the participation ratio (PR) (***Abbott et al., 2011***; ***Mazzucato and La Camera, 2016***; ***Gao et al., 2017***). The PR depends on all covariances in the network as it is derived from the eigenvalues of the covariance matrix using a principle component analysis. One can show that it depends on a combination of first and second order moments of auto- and cross-covariances (***Mazzucato and La Camera, 2016***). The physical interpretation of the PR is the dimensionality of the manifold spanned by the neuronal activity (see Sec. Materials and Methods: Covariances and dimensionality). The higher the PR, the more eigenvectors (principle components) are needed to capture most of the variance of the dynamics. We performed an analysis for the REST and also for the R2G states (0.5 s slices were concatenated to 3 s slices). To make the PR of different experiments and recordings comparable, we normalized to the total number of SU obtained in each session.

Figure 4B shows that the dimensionality varies over time (shown for monkey E during REST experiment). It changes with relation to behavior consistently across monkeys (as shown in Fig. 4C). This is true for all sessions (see also Tab. 6). The PR is highest during RS and PP and lowest during TM. The RSS state in both monkeys is clearly distinct from RS, its PR being more similar to the one obtained for M, as seen in the covariance distributions. The spread of the values is notably higher in REST than in R2G states, especially in monkey N.

**Table 6.** Quantification and correlation of balance and dimensionality. Top rows: quantification of balance between putative excitatory and inhibitory population activities (Δ(bs, ns) and *ρ*(bs, ns)). Middle row: quantification of dimensionality as measured by the participation ratio (PR). Bottom row: Spearman rank correlation between *ρ*(bs, ns) and PR; only *p*-values smaller than 0.05 are listed. All values were obtained from 3 s slices, for different behavioral states in REST (RSS, RS, M) and R2G (TM, PP), pooled across all recordings of the respective type.

| Monkey | RSS | RS | M | TM | PP |
| --- | --- | --- | --- | --- | --- |
| Putative excitatory/inhibitory prevalence $\Delta(\text{bs, ns})$ | | | | | |
| E | $-0.15 \pm 0.86$ | $0.00 \pm 0.72$ | $0.06 \pm 0.85$ | $-0.16 \pm 1.04$ | $0.16 \pm 0.89$ |
| N | $-0.32 \pm 1.00$ | $0.09 \pm 0.98$ | $-0.06 \pm 1.07$ | $-0.28 \pm 0.97$ | $0.28 \pm 0.68$ |
| Instantaneous balance $\rho(\text{bs, ns})$ | | | | | |
| E | $0.39 \pm 0.22$ | $0.2 \pm 0.23$ | $0.44 \pm 0.29$ | $0.26 \pm 0.22$ | $0.1 \pm 0.21$ |
| N | $0.36 \pm 0.21$ | $0.17 \pm 0.24$ | $0.16 \pm 0.24$ | $0.39 \pm 0.24$ | $0.08 \pm 0.21$ |
| Participation ratio PR |  |  |  |  |  |
| E | $0.100 \pm 0.015$ | $0.135 \pm 0.019$ | $0.111 \pm 0.028$ | $0.097 \pm 0.015$ | $0.152 \pm 0.018$ |
| N | $0.101 \pm 0.018$ | $0.130 \pm 0.023$ | $0.111 \pm 0.023$ | $0.071 \pm 0.012$ | $0.134 \pm 0.010$ |
| Spearman rank correlation between $\rho(\text{bs, ns})$ and PR | | | | | |
| E | -0.31 | $-0.44 (p \ll 0.001)$ | $-0.77 (p \ll 0.001)$ | 0.14 | 0.04 |
| N | $-0.32 (p < 0.01)$ | $-0.21 (p < 0.001)$ | $-0.42 (p < 0.0001)$ | $-0.33 (p < 0.001)$ | -0.08 |

Kruskal-Wallis tests on all behavioral conditions yield highly significant differences (*p*≪ 0.0001) for both monkeys. Pairwise comparisons yield mostly significant results except for RS vs PP, RSS vs M & TM and M vs TM in monkey E, as well as RS vs PP and RSS vs M in monkey N. These results hold for different bin sizes (shown in Sec. Materials and Methods: Covariances and dimensionality).

The higher dimensionality of RS as compared to movement states and sleepy rest is a clear evidence for the complexity of this state. Moreover, the large difference between the PR of RS and RSS emphasizes the necessity to distinguish between rest with eyes open and closed. In the following, we will support this claim by analyzing the balance between putative excitatory and inhibitory population activity.

### Balance in population activity

Population activities are the most straight forward and well studied low-dimensional projections of neuronal spiking data. Based on summed SU activities, they provide a global view on the network activity, disregarding single neuron-specific fluctuations. Due to the population-averaging, fluctuations on the population level are only determined by the average single-neuron covariances (***Kriener et al., 2008***) and are insensitive to the large variability across single neurons (Fig. 4A). The latter, in contrast, affects the participation ratio (***Mazzucato and La Camera, 2016***). Studying population-level coordination is therefore complementary to the analysis of dimensionality.

Balance between excitation and inhibition is considered an attribute of a physiological network state in contrast to non-physiological states like, e.g., epilepsy (***Zhang and Sun, 2011***; ***Dehghani et al., 2016***). Theoretical studies simulating cortical network dynamics mostly assume a balanced resting state (***van Vreeswijk and Sompolinsky, 1996, 1998***; ***Brunel, 2000***) and relate this to low average covariances between neurons (***Renart et al., 2010***; ***Tetzlaff et al., 2012***). We here investigate the balance between putative excitatory (bs) and inhibitory (ns) population activities, first globally, similar to ***Dehghani et al. (2016)***, and then relating the balance levels to different behavioral states. Figure 5A shows the deviations from balance at different time scales (bin width) in session E1.

**Figure 5.**
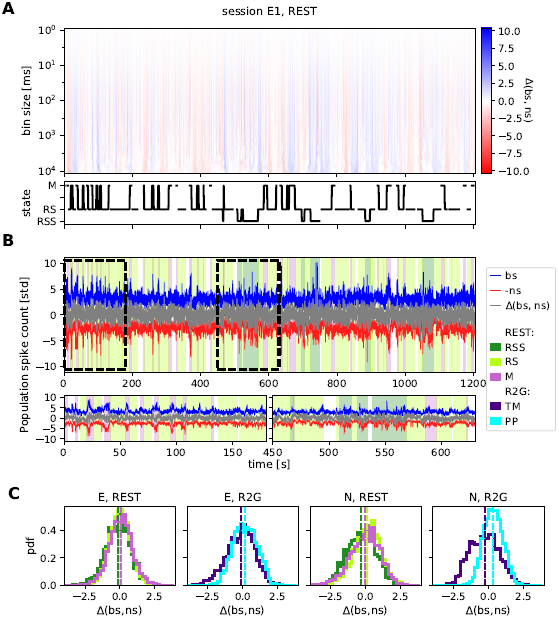
Balance between putative excitatory and inhibitory population activity. **(A)** Multiscale balance during a single REST session of monkey E. The x-axis (shared with B) indicates time, the y-axis indicates the temporal resolution (bin size), the color marks the difference between z-scored putative excitatory and inhibitory population activities. The black trace below indicates behavioral states (cf. Fig. 2A). **(B)** Close-up view on a 100 ms scale. Population activities and the difference (grey) between z-scored putative excitatory and inhibitory firing from the same session as in A. Colors in the background denote behavioral states (cf. Fig. 1A). For a better visualization, spike counts of bs (blue) and ns (red) populations, calculated in 100 ms bins, are normalized by their standard deviation instead of z-scoring. Additionally, ns time series is multiplied by (−1). **(C)** Histograms of the difference between globally z-scored population activities of putative excitatory (bs) and inhibitory (ns) SUs of all REST (left) and R2G sessions (right) for monkey E (first two panels) and monkey N (last two panels), calculated in 100 ms bins. Results are pooled across all recordings of the respective type.

White color indicates values close to zero, i.e., well-balanced activity, prevalent on smaller time scales. On time scales larger than ≈30 ms, blue and red vertical stripes indicate transient deviations from perfect balance, i.e., an instantaneous dominance of excitation or inhibition, respectively. Such brief fluctuations were also observed during physiological activity by ***Dehghani et al. (2016)***.

Fig. 5B presents a detailed view on one single time scale: the spike counts in 100 ms bins of bs (blue) and ns (red) population and the difference between z-scored population spike counts in grey, representing a horizontal slice of Fig. 5A. Putative excitatory and inhibitory activities seem to fluctuate simultaneously, indicating balance, although the considered bin size is much larger than 30 ms: Pronounced deviations from average spike counts can be seen in both populations, especially during RSS (dark green background color) and M (pink background). Considering the distributions of mean population spike counts (Appendix 1 Fig. 1), the standard deviations during M and RSS (9.56±2.49 and 7.81±2.75, respectively, ns population, session E1) are much higher than during RS (7.56±1.54). They are even larger (approximately factor 1.7) than expected from the larger means (approximately factor 1.1) which indicates distributions with more extreme values, i.e., potential transient increases in the population spike count.

We performed a quantitative analysis of how the balance between bs and ns SUs relates to the behavioral states, on the time scale of 100 ms, for our REST and R2G data.

Firstly, we asked if there was a state-specific prevalence of ns or bs activity. Fig. 5C shows the results of subtracting z-scored ns from z-scored bs population activity. The histograms show the distributions of values obtained in all 100 ms bins in the pooled REST and R2G sessions of monkey E and N, and Tab. 6 (top row) lists the values of mean and standard deviation.

We find a clear shift between PP and TM distributions in the R2G data of both monkeys: TM distributions are shifted towards negative and PP towards positive difference values (mean±std: 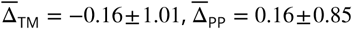 for monkey E and 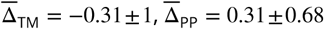 for monkey N), pointing out a prevalence of ns or bs activity, respectively. This indicates that the balance between the excitatory and inhibitory activity dynamically changes depending on the behavioral state of the monkey during task performance.

In the REST data of both monkeys, the RSS state is dominated by inhibition (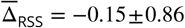 for monkey E and 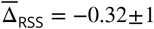 for monkey N). Concerning RS and M, however, the general tendencies are less pronounced and inconsistent: For monkey E, we find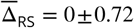 and 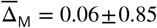, thus no particular dominance. For monkey N, we find a slight tendency of movements being dominated by putative inhibitory firing similarly to 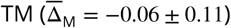, while resting periods are again not significantly dominated by any population 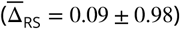.

Secondly, we quantified the level of instantaneous balance by computing the Spearman rank correlation *ρ*(bs, ns) between bs and ns population activity in 3 s data slices (cf. ***Renart et al. (2010)***; ***Tetzlaff et al. (2012)***). A higher correlation value indicates a more strict instantaneous balancing between the excitatory and inhibitory activity. Fig. 6A and B show box plots of the correlation measure *ρ*(bs, ns) for the different behavioral states of the two monkeys, and the corresponding means and standard deviations are listed in Tab. 6. For monkey E, the correlation between bs and ns activity is highest during 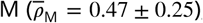, meaning that the balance was kept best during M state, closely followed by 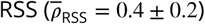, see Fig. 6A, left. RS shows the lowest correlation 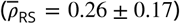, it is thus the least balanced state during REST. Pairwise comparisons confirm significantly different results for RS vs M, but not for RSS vs M & RS. In monkey N, RS and M exhibit nearly identical correlations 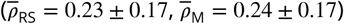, see Fig. 6B, both are less balanced than 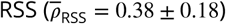 which is significantly more balanced than M.

**Figure 6.**
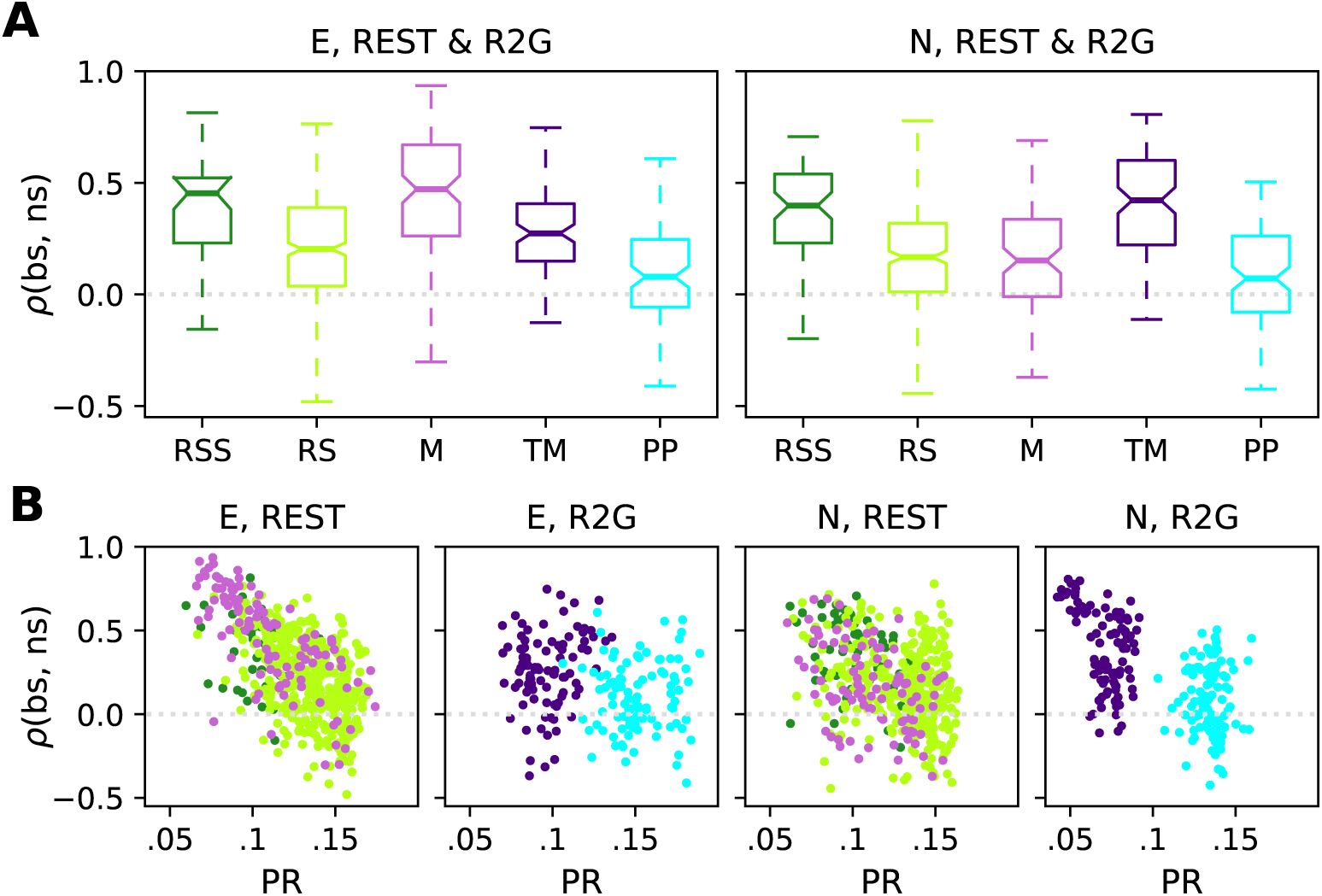
Instantaneous balance and its relation to dimensionality. **(A)** Box plots of the correlation between putative excitatory and inhibitory population activity calculated in 3 s slices, which quantifies the instantaneous (100 ms) balance for monkey E (left) and N (right). **(B)** Scatter plots showing the relationship between the instantaneous balance *ρ*(bs, ns) and dimensionality PR, for monkey E (panels on the left) and N (panels on the right). Each dot represents the PR and *ρ*(bs, ns) values of one 3 s slice during REST or R2G recording. Results are pooled across all recordings of the respective type.

In the R2G data of both monkeys (right panels of Fig. 6A and B, respectively), PP (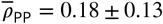 for monkey E and 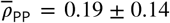 for monkey N) is less balanced than TM (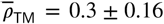 for monkey E and 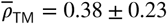 for monkey N); PP shows a significantly (*p<* 0.001) lower correlation between ns and bs activities. We thus conclude that behavioral states without movements (RS, PP) are less balanced than movement states when considering a timescale of 100 ms.

Participation ratio and balance measure different aspects of correlations in the underlying network. We now ask if and how these measures relate to each other. To this end, we analyzed the relation of PR and *ρ*(bs, ns) using scatter plots (Fig. 8C & D)—each 3s slice is represented by a single data point. The points are colored according to the behavioral state they are computed from; RSS, RS & M for REST, and TM & PP during R2G. For the REST data, we observe a negative correlation between PR and *ρ*(bs, ns) (see Tab.6): The higher the complexity, the lower the balance. Data points from different behavioral states overlap strongly and are thus not clearly separable. In contrast, TM and PP of the R2G data separate into two different clouds according to their PR, but there is no clear correlation to *ρ*(bs, ns).

**Figure 7.**
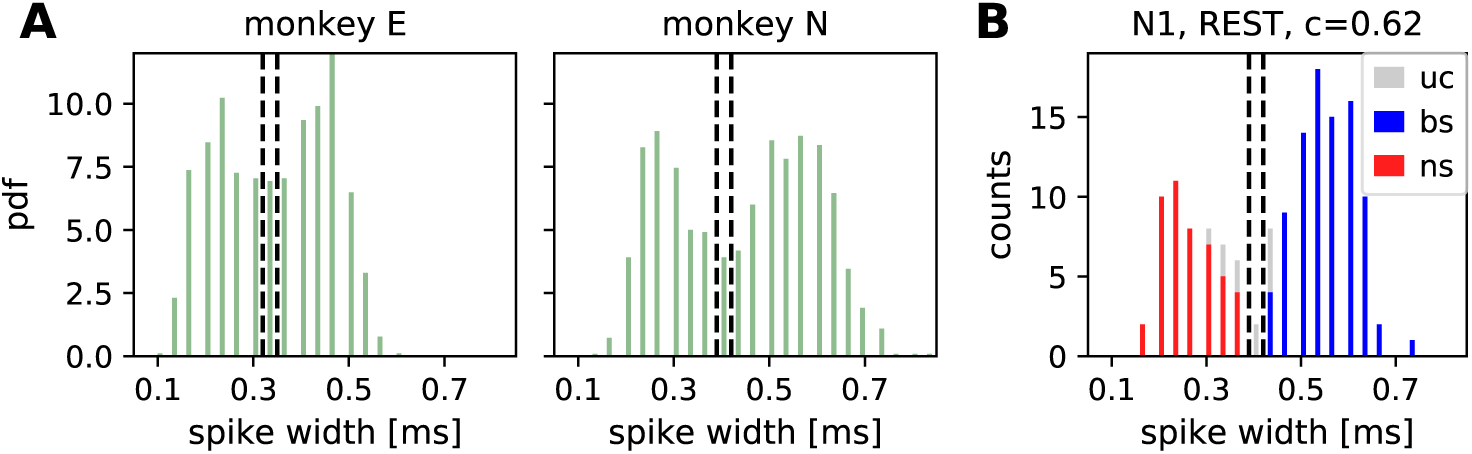
Separation between broad-spiking (bs) and narrow-spiking (ns) single units (SUs). **(A)** Spike-width distributions calculated from the average spike widths of all REST and R2G recording sessions for monkey E and monkey N. The two vertical lines indicate the thresholds for ns (0.33 ms and 0.4 ms for monkeys E and N, respectively) and bs units (0.34 ms and 0.41 ms for monkeys E and N, respectively). **(B)** Exemplary separation between ns (red) and bs (blue) units for the first REST recording of monkey N (N1). SUs between the thresholds are left unclassified (grey), as well as all SUs with a consistency smaller than 62%.

**Figure 8.**
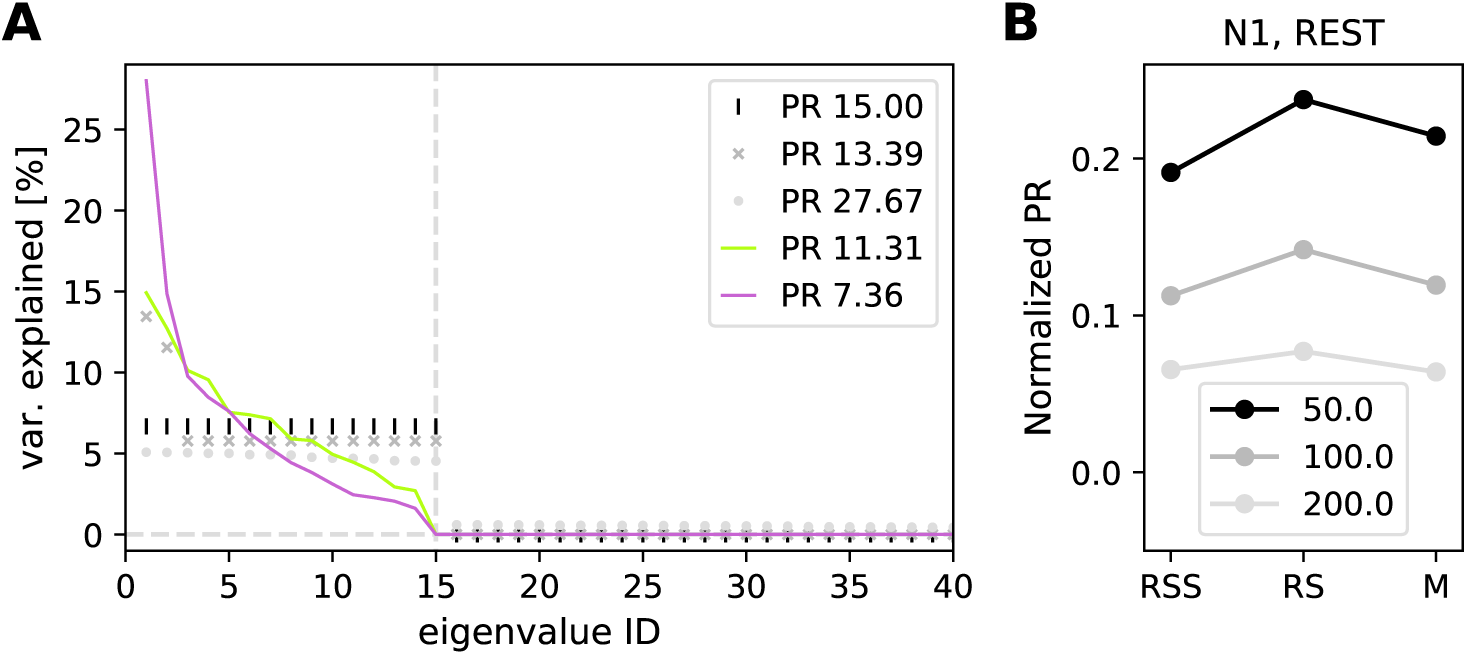
Participation ratio to characterize the dimensionality. **(A)** Sketch showing relation between the eigenvalue spectrum and PR. In case of first *N* eigenvalues explaining equal amounts of variance and the rest vanishing, the PR equals *N* (black vertical lines). If a few eigenvalues are much higher than the others, the resulting PR decreases (dark grey crosses). If, on the contrary, a uniformly distributed random value is added to each eigenvalue from the first case, the calculated PR becomes higher (light grey circles). Experimental data is typically a mixture of the second and the third case. Continuous traces show exemplary results for a single 3 s data slice of RS (green) and M (pink) in session N1. **(B)** Participation ratio, calculated with different bin sizes (50, 100, 200 ms) in exemplary session N1.

## Discussion

Experiments without any imposed stimuli or task have been investigated in numerous studies and referred to with multiple names: (a) ongoing, intrinsic or baseline activity of single brain areas (***Arieli et al., 1996***; ***Tsodyks et al., 1999)***, (b) spontaneous or resting state activity on the whole brain level (***Vincent et al., 2007***; ***Raichle, 2009***; ***Deco et al., 2011)***, as well as (c) idle state of point-neuron network simulations (***Brunel, 2000***; ***Potjans and Diesmann, 2014***; ***Dahmen et al., 2019)***. Yet, a thorough characterization of spiking activity in the resting condition on the level of single neurons was still missing.

Here, we investigate the properties of spiking activity in macaque motor cortex during five behavioral states: resting state (no movements, RS), sleepy rest (no movements with eyes closed, RSS), spontaneous movement (M), task-related movement (TM) and task-imposed waiting without movements (PP), with a particular focus on RS. Our main findings are: (a) we demonstrate a considerable correlation between neuronal firing and behavior, (b) we find that RS single unit activity is characterized by relatively low average firing rates and a high variability of interspike intervals and spike counts across data slices, (c) we identify a high dimensionality of the joint activity during RS, which is (d) correlated with a low level of balance between putative excitatory and inhibitory population spiking.

### Single unit activity and LFP during different behaviors

Many studies investigate the link between neuronal activity in the motor cortex and behavior using LFP data (e.g. ***Pfurtscheller and Aranibar (1979)***; ***Fontanini and Katz (2008)***; ***Engel and Fries (2010)***; ***Kilavik et al. (2013)***). Low frequency oscillations (<15 Hz) are often linked to sleep (***Gervasoni et al., 2004***; ***Fontanini and Katz, 2008)***, beta oscillations (≈13-30 Hz) typically appear during movement preparation or postural maintenance (***Baker et al., 1999***; ***Kilavik et al., 2012)***, while faster oscillations mostly reflect attention and neuronal processing during movements (***Fontanini and Katz, 2008***; ***Liu and Newsome, 2006)***. Our visual classification of the behavior is in good agreement with the LFP characteristics shown in the above studies.

Firstly, all states without movements (RSS, RS & PP) show pronounced beta oscillations which are shifted towards higher frequencies during task-imposed rest (PP) compared to spontaneous rest (RS). Secondly, both spontaneous and task-related movements (M & TM) show stronger fast oscillations than non-movement states. The spectra obtained during RSS (eyes closed) indicate distinct physiological states in the two monkeys: the peak frequency during RSS of monkey E occurs at a much lower frequency compared to monkey N. This suggests that closing the eyes indicates drowsiness in monkey E but not necessarily in monkey N.

Furthermore, in agreement with previous studies on behaving monkeys (***Nawrot et al., 2008***; ***Nawrot, 2010***; ***Rickert et al., 2009***; ***Churchland et al., 2010***; ***Riehle et al., 2018)***, we find that the spiking activity is highly variable across SUs, and that the average firing rate is increased during movements as compared to waiting for the cue at rest. In REST data, in ≈50% of all SUs, we find a significant correlation between SU firing and the monkey’s behavior. This indicates that the analysis of spiking activity is another valid approach next to LFP and large scale recordings to investigate behavioral states, including resting state. Analogously to activations and deactivations of specific brain areas reported in fMRI studies (***Biswal et al., 1995***; ***Raichle, 2009***; ***Deco et al., 2011)***, we observe systematic in- and decreases in firing rates in numerous SUs. Also in agreement with ***Nawrot et al. (2008)***; ***Nawrot (2010)***; ***Rickert et al. (2009)***; ***Churchland et al. (2010)***; ***Riehle et al. (2018)***, we find a (slightly) lower spike count variability during task related movements (TM) than during movement preparation (PP) and vice versa for the spike time irregularity^1^.

A new finding of our study is a pronounced difference in variability between REST and R2G states, i.e., between spontaneous and task-related behavior. All REST states show a significantly higher spike count variability and a higher firing irregularity than the R2G states. These differences are probably due to the behavioral constraints present in the R2G but not in the REST experiments. During R2G task, the monkey received visual input to control periods of waiting or arm movements, resulting in well-defined behavioral states and partially constrained mental states with a more regular and reliable firing. In contrast, during REST experiments, the monkey itself decided what to do (e.g. movement preparation or onset), resulting in a less well-defined behavior and its timing.

The above findings are consistent for the two monkeys, but there are differences concerning the sleepy resting state: For monkey E, firing rates during RSS are higher than during RS, thus closer to the values measured during M, while this is not the case for monkey N. Thus, similar to what we find for the LFP spectra, the distinction between RS and RSS (eyes open vs eyes closed) is more pronounced in monkey E than in monkey N.

### Network activity

During all behavioral states, the network activity of groups of neurons in the motor cortex is characterized by a dimensionality much lower than the maximal possible dimension, i.e., the total number of recorded single neurons. Task-related movements show the lowest dimensionality, expressed by a small normalized participation ratio, while non-movement states show a higher dimensionality. Accordingly, neuronal firing during rest is less coordinated than during other states, as indicated already by the narrower covariance distribution centered at zero. These findings agree well with ***Mazzucato and La Camera (2016)***; ***Gao et al. (2017)*** who compare stimulus-evoked and ongoing neuronal activity, assuming M and TM to represent the evoked activity, and RS and PP the ongoing activity. The low normalized participation ratio of less than 0.1 during TM (Figure 4) shows that the neural state space dynamics of the reach-to-grasp movement can be reconstructed from only a few principal components. Given the number of observed SUs (see Table 1), this corresponds to a neural state space dimensionality of approximately 7-13. In contrast, the ongoing activity during RS and PP is of significantly higher dimensionality (≈12-17 and ≈12-22, respectively) and thus more complex.

In accordance with ***Csicsvari et al. (1999)***; ***Peyrache et al. (2012)***; ***Dehghani et al. (2016)***, we also find that putative excitatory and inhibitory population spiking are primarily well balanced. However, our detailed time-resolved analysis, i.e., calculating the balance in 100 ms bins, uncovers the following particularities. During R2G experiments, the activity alternates between excitation-dominated movement preparation (PP) and inhibition-dominated movement execution (TM). During non-movement states (PP and RS), we find a reduced correlation between putative excitation and inhibition, i.e., a reduced instantaneous balance of non-movement states. In addition, the instantaneous balance is anti-correlated to the dimensionality, particularly strongly in REST.

We suspect that the relatively high instantaneous balance during movements and sleepy rest is partially an effect of an enhanced number of transient changes in population spiking in these states as compared to the other states (Fig. 5B). A prominent increase in firing as observed during movements is an unambiguous type of activity change and is thus easy to capture by correlation measures. Such transient increases correlated in time between two neuronal populations could result from the recurrent coupling between excitatory and inhibitory neurons (see Appendix 1). In addition to the transients in population activity, we find hints of a prevalence of non-stationarities (e.g. transients) in the SU firing during movements and sleepiness, but not during rest (Appendix 1 Fig. 2). Strong transient comodulations of spiking activities amplify correlations between neurons, which in turn decrease the dimensionality of network activity. Therefore, transient changes in firing rates might also be partially responsible for the reduced dimensionality during task-related movements.

### Influence of pre-processing and critical assumptions

Specificities of extracellular recordings and the following pre-processing steps impose particular biases on the resulting statistics and their interpretation. Firstly, the spike sorting procedure, necessary to identify single cells recorded on the same electrode, is well-known to be problematic (***Lewicki, 1998***; ***Quian Quiroga, 2012)***. Additional limitations on minimal SNR and firing rate of a sorted unit to be considered for statistical evaluation contributes to the undersampling of sparsely firing neurons and thus biases results towards highly active neurons. This is often referred to as the problem of “dark matter” of the brain (***Shoham et al., 2006)***.

Secondly, the separation between putative excitatory and inhibitory neurons based on the widths of their spike waveforms is also known to have several limitations (***Bartho et al., 2004***; ***Kaufman et al., 2010, 2013***; ***Peyrache et al., 2012***; ***Dehghani et al., 2016***; ***Peyrache and Destexhe, 2019)***. Some pyramidal neurons, in particular when recorded close to the axon, exhibit narrow waveforms. Still, it was shown that over 10% of M1 interneurons have intermediate or broad waveforms (***Kaufman et al., 2010***; ***Vigneswaran et al., 2011***; ***Kaufman et al., 2013)***. When discussing the differences between the two populations, it should be kept in mind that not all narrow-spiking units are inhibitory and only a part (majority) of broad-spiking SUs are excitatory (***Peyrache and Destexhe, 2019)***. Nevertheless, our separation yields higher average firing rates for putative inhibitory neurons which agrees well with what is known from the literature (***Peyrache et al., 2012***; ***Dehghani et al., 2016***; ***Kaufman et al., 2010)***.

Thirdly, our study relies on the behavioral segmentation of REST recordings which is highly subjective and has rather poor temporal resolution (∼1 s) in comparison to the recorded neuronal activity (∼1 ms). Nevertheless, our behavioral classification seems to be accurate in terms of separating sets of dissimilar neurophysiological network states, as reflected by differences in state-resolved LFP spectra, see above. Still, our definitions of the behavioral states are based on visual inspection and may not be as precise. For example, the identification of “whole body and limb movements” in the video recording does not account for the fact that, due to the exact placement of the Utah array, our recordings are particularly sensitive to contra-lateral arm movements. Likewise, the RS classification is simply based on the exclusion of movements with the additional criterion of “eyes open". Compared to the very precise behavioral classification in R2G recordings^2^, the behavioral segmentation of REST recordings is vague and allows for a much broader range of actual behaviors.

Finally, reliable covariance estimation necessitates very long data slices (***Cohen and Kohn, 2011)***. To satisfy this requirement, in R2G data we had to concatenate slices from 6 consecutive trials into 3 s slices for the analysis of covariance and participation ratio. Thus, a single PR value results from averaging over six independent recording periods in contrast to the continuous REST data. However, this approach can be justified by our observation of a low inter-trial variability obtained for 0.5 s slices of the R2G data.

### Towards experimental data for spiking model validation

Modeling studies focusing on spiking-neuron networks often claim to model an “idle” state, i.e. without any relation to functional aspects, characterized by sparse asynchronous irregular spiking and balanced input statistics (***van Vreeswijk and Sompolinsky, 1996***; ***Amit and Brunel, 1997***; ***van Vreeswijk and Sompolinsky, 1998***; ***Brunel, 2000***; ***Kumar et al., 2008***; ***Voges and Perrinet, 2010, 2012***; ***Potjans and Diesmann, 2014)***. To isolate the ongoing and recurrently generated activity, many of these studies consider stationary states without any transient network activation due to external inputs. In this case single-neuron and population firing rates fluctuate around some mean activity. However, data collected in behavioral experiments often contain transient firing rate fluctuations on the level of both single units and whole populations. For motor cortex recordings, such firing rate changes typically occur during movements, which has been shown here and in many other studies (***Nawrot et al., 2008***; ***Rickert et al., 2009***; ***Churchland et al., 2010***; ***Riehle et al., 2013, 2018)***. We find that this disagreement can (mostly) be avoided by considering resting periods (RS) in REST recordings only. Using non-movement epochs (PP) during behavioral tasks yields results that are more similar to RS in terms of network firing properties, but the SU variability is still different (higher for CV2, and lower for FF). A comparison to inappropriate data sets could lead to erroneous conclusions on model parameters and the mechanisms that shape the network dynamics. Hence, network models that claim to mimic an idle state in terms of SU and network activity should ideally be validated against resting state data.

#### Balance and correlations

Another typical claim of network simulations is the assumption of a balanced state, see above. The modeling literature discusses different types of balance (***Deneve and Machens, 2016)***. Many studies assume a cancellation of excitation and inhibition in the *input* to neurons based on a balance between the strength and number of excitatory and inhibitory afferent connections (***Poil et al., 2012)***. Perfect balance in this context corresponds to a critical point, where network dynamics exhibits avalanche-like behavior (***Beggs and Plenz, 2003)***. This static notion of balance purely relies on the network structure. In contrast, other studies describe a more or less tight “dynamical balance” (***van Vreeswijk and Sompolinsky, 1996***; ***Amit and Brunel, 1997***; ***van Vreeswijk and Sompolinsky, 1998***; ***Brunel, 2000)***, where excitatory and inhibitory inputs cancel each other at each point in time (***Renart et al., 2010)***. The latter cancellation is caused by excess inhibitory feedback (***Tetzlaff et al., 2012)*** and, in excitatory-inhibitory networks, is accompanied by correlations between excitatory and inhibitory spiking (***Renart et al., 2010)***. These correlations can be quantified on the level of neuronal *output*. Therefore, we here study balance based on the correlation between population activities.

Our observation of a reduced instantaneous balance during resting state compared to other states at first sight seems to argue against model validation with RS data. However, the balanced state does not necessarily rely on instant tracking between excitatory and inhibitory population activities. What it demands instead is a cancellation in the input to each single neuron, which does not uniquely define a correlation structure between outputs (***Helias et al., 2014)***. Deviations between population activities can indeed be organized such that their net effect to the summed input to single neurons cancels out (***Tetzlaff et al., 2012***; ***Baker et al., 2019)***. Furthermore, one should keep in mind that we investigate a rather large time scale, and the apparent reduction of balance could be an effect of fewer transient activities contributing to the correlation between excitatory and inhibitory population activities during rest. Nevertheless, we find principally well-balanced population firing in all behavioral states and we show that spiking during rest is neither dominated by excitation nor by inhibition which indicates that RS periods are in agreement with balanced network models.

Related to balance, modelers often assume uncorrelated or weakly correlated external inputs to local networks, but it is impossible to determine the amount of correlations in the neuronal input with extracellular recordings. Strongly correlated inputs, attributed to sensory (***Decharms and Merzenich, 1996)*** or movement processing (***Murphy et al., 1985)***, may boost the modulation of firing rates on the population level. This could lead to higher pairwise covariances and subsequently lower dimensionalities than expected in artificial networks with a well controlled input structure. We find that such a decrease in dimensionality, is, for example, particularly pronounced during task-induced movements. This again points out the necessity to separate between rest and movements in order to avoid potential unrealistic mismatch between input and output statistics of spiking models.

#### Heterogeneity of neuronal networks

Another point is the remarkable heterogeneity of neuronal activities in experimental recordings: SUs show a broad range of firing rate profiles and spiking (ir-)regularities, as well as distinct activity modulation related to behavioral state changes. Neuronal network studies mostly are able to reproduce this heterogeneity. Single-neuron properties (e.g. time constants, synaptic weights) and connectivities are typically given as parameter ranges described by certain distributions derived from experimental measurements (***Kumar et al., 2008***; ***Voges and Perrinet, 2012***; ***Potjans and Diesmann, 2014)***. Depending on the widths of these distributions (and other features) the resulting activities can and should be adapted to the heterogeneity in experimental data (***Dahmen et al., 2019)***. An advantage of heterogeneous network activity is that it enhances the stability of the “idle” state (***Denker et al., 2004)*** which is essential for real-world neuronal networks that need to be able to operate under various conditions. For example, the different behavioral states analyzed here demonstrate that the motor cortex operates in similar dynamical regimes for various kinds of behaviors, including movements and sleepiness. The stability range of network models can be further increased by including more real-world features like homeostatic mechanisms (e.g. adaptation, short term plasticity) which also support a high (temporal) heterogeneity.

In summary, we encourage modelers to (continue to) incorporate the heterogeneity of real-world neuronal activities and we conclude that the validation of network models that claim to simulate idle states should be based on resting state data. Still, even when considering REST recordings without any task or stimulus, it is necessary to separate out the “pure” resting state periods because they show distinct statistical properties: lower firing rates, fewer transient activities, smaller covariances and thus a higher dimensionality.

### Definition of behavioral states

The rather vague classification of behavioral states in REST recordings is based on observing the monkey in contrast to the precise classification in R2G experiments which relies on external cues. The consequence of this difference in precision is clearly visible on the level of the spiking activity statistics: In addition to the higher spike time irregularity and the higher (broadly distributed) spike count variability in REST compared to R2G, REST states also show a less clear state-specific difference in the dimensionality results.

In addition, there is the problem of different time scales (i.e., slice lengths): 0.5 s as forced by the R2G settings versus the heuristically chosen 3 s in REST. Thus, some comparisons between single behavioral states of these two data types might be unfair, but we still observe the expected commonalities in the states with (TM, M) versus without movements (PP, RS): Non-movement states show generally lower firing rates, a higher dimensionality, and a lower instantaneous balance.

#### Eyes open vs closed

The sleepy resting state RSS, however, turns out to be a special case. As already mentioned, the LFP spectra during RSS and the firing statistics of RSS are monkey-specific: in monkey E, the distinction between RS and RSS is more pronounced than in monkey N. However, concerning both dimensionality and instantaneous balance, the RSS distributions of the monkeys are similar. In addition, mean dimensionalities are closer to the ones obtained for M than for RS, even though RSS is a non-moving state. In accordance with observations that the motor cortex can show distinct reactions to visual stimuli (***Wannier et al., 1989***; ***Riehle, 1991)***, we conclude that the distinction between eyes-open and eyes-closed is important even in the motor cortex, since there is an impact on the neuronal activity. RS and RSS can be distinct physiological states in a given monkey: monkey E seems to be really drowsy when its eyes are closed while monkey N might be simply bored. This example also shows the importance of verifying the result of the visual behavioral segmentation with the LFP spectra of the resulting states.

#### Alternative classification methods

There are other possibilities for the behavioral segmentation of REST recordings. One idea would be an automatic decoding of behavioral state purely based on SU firing properties by means of machine learning methods, e.g., ***Pandarinath et al. (2018)***. Given that approximately 50% of all SUs exhibit a strong correlation between firing rate modulations and behavior, such an approach would probably be possible but not necessarily straight-forward. If there were enough data to define an appropriate learn set, a machine learning algorithm could, for example, identify SUs that consistently increase or decrease their firing rate with specific state changes. Such an approach, however, is beyond the scope of this study. Another idea would be to increase the temporal precision of the visual segmentation by means of an automated detection of transient neuronal activities. Yet, the detection of transient activities in itself is not trivial (***Ito et al., 2019)***, it does not allow to distinguish between RSS and M, and particularly in our data a 3 s long movement epoch contains several such transients in an unknown frequency. We do not pursue this approach, as it is again beyond the scope of this study.

#### Resting state as superposition of sub-network activities

An interesting hypothesis emerges from the comparison of our study to resting state studies based on large-scale measurements. Similar to the observation of activations and deactivations of specific brain areas in fMRI studies (***Biswal et al., 1995***; ***Fox and Raichle, 2007***; ***Raichle, 2009***; ***van den Heuvel and Hulshoff Pol, 2010***; ***Deco et al., 2011)***, we observe systematic in- and decreases in the spiking activity of numerous SUs. Large-scale studies conclude that spontaneous brain activity emerges from a set of resting state networks (***Fox and Raichle, 2007***; ***Raichle, 2009***; ***van den Heuvel and Hulshoff Pol, 2010***; ***Deco et al., 2011)***, i.e., from a sequence of consistently re-occurring spatio-temporal activity patterns that resemble task-evoked activity, but are present during rest (***Vincent et al., 2007***; ***Fox and Raichle, 2007***; ***van den Heuvel and Hulshoff Pol, 2010)***. One could thus hypothesize a similar phenomenon on the microscopic level of spiking activity: a resting state composed of the activities of several sub-networks of single neurons in the motor cortex. During movements, one could imagine a convergence of the neuronal activity into specific networks (cf. (***Fox and Raichle, 2007***; ***Mazzucato and La Camera, 2016)***). The larger spatial spread of the activity observed during RS compared to M (see Appendix 2) would be in line with the above hypothesis, assuming that a superposition of many spatially embedded networks yields an enlarged spatial extent than a single such network (cf. Fig. 1 in Appendix 2). Likewise, the high dimensionality observed during RS agrees well with the hypothesis of a superposition of several sub-networks.

Yet another question concerns the definition of “rest” in general: how to define it in other cortical areas than motor cortex, e.g., in sensory systems? For the auditory system one would intuitively assume that silence or white noise as auditory input represents the resting condition. Similarly, for the visual system one could use a uniform or noise background as visual input. The choice of “eyes-closed” as rest condition would, however, represent a different behavioral state compared to our assumption of sleepy rest being a qualitatively different condition.

Given all the issues concerning the definition of “rest” and the behavioral segmentation, together with the superposition of RSNs on the scale of brain areas, one could claim that it is futile to attempt to characterize the spiking activity during an assumed resting state. However, our results clearly demonstrate a set of significant differences between the spiking activity in motor cortex during “rest” as compared to other behavioral conditions.

### Conclusions

We demonstrate that spiking activity in monkey motor cortex during rest differs significantly from other spontaneous and task-related behavioral states, for example sleepiness and movements. The main characteristics of the resting state activity are low average firing rates combined with a high variability of single-unit spiking statistics, and a pronounced complexity as indicated by a less coordinated firing which results in a higher dimensionality of the network activity. We show that and explain why neuronal network models should be validated against resting state data, aiming to enhance the trend towards realistic network models that account for the heterogeneity in neuronal data. We hope that our study is just the beginning of the characterization of “rest” on the level of spiking neurons. More specific analysis is needed to quantify transient activities, their relation to the balance between exitatory and inhibitory population activities, and to provide an automated algorithm for the behavioral segmentation of REST recordings.

## Materials and Methods

We first describe the two types of experimental recordings analyzed in this paper: resting state (REST) and reach-to-grasp (R2G) data, the latter obtained during a behavioral task. Then, we explain the experimental procedure and the pre-processing of all data types with a particular focus on the REST recordings and their behavioral classification. Finally, the measures calculated to characterize different behavioral states are listed and explained.

### Experimental paradigm and recordings

Two adult macaque monkeys (*Macaca mulatta*), female (monkey E) and male (monkey N), participated in two distinct behavioral experiments: resting state (REST) and reach-to-grasp (R2G). Monkeys were chronically implanted with a 4×4 mm^2^ 100 electrode Utah Array (Blackrock Microsystems, Salt Lake City, UT, USA) situated in the hand-movement area of (pre-)motor cortex. Spiking activity and LFP were recorded continuously during an experimental session, with sampling frequency of 30 kHz. Details on surgery, recordings and spike sorting, as well as on the R2G settings are described in ***Riehle et al. (2013, 2018)***; ***Brochier et al. (2018)***

During a resting state session, the monkey was seated (but not fixated) in a primate chair. The chair was positioned so as to prohibit the animal from reaching any objects. There was neither a particular stimulus nor any task, the monkey was free to look around and move spontaneously. In addition to the registration of brain activity, the monkey’s behavior was video recorded and synchronized with the electrophysiology. For each monkey two such sessions were recorded and lasted approximately 15 min for monkey N and 20 min for monkey E.

In the R2G experiments, monkeys were trained to perform an instructed delayed reach-to-grasp task to obtain a reward, see Fig. 1B. The monkey had to self-initiate a trial by closing a switch (TS). After 800 ms a CUE-ON signal provided some task-related information. 300 ms later, the CUE signal was switched off which defined the start of the preparatory period, during which the monkey was supposed to sit still. One second after the CUE-OFF, a GO signal provided the complementary task-related information and indicated the monkey to start moving. The monkey had to release the switch (SR) and reach to the target. After grasping the object, the monkey had to pull and hold it for 500 ms to obtain the reward (RW). Brain activity was recorded together with time stamps of all events within a trial.

Table 1 lists all single recording sessions for both REST and R2G experiments. Typically, a REST recording was performed subsequent to an R2G recording session. Only the E2 session was recorded directly before an R2G session which is probably the reason for the missing RSS intervals. The monkey was rather twitchy, impatiently waiting for the R2G tasks, because R2G experiments include a reward while there was no reward during REST recordings.

### Pre-processing

The waveforms of potential spikes were sorted into the SUs offline and separately on each electrode using the Plexon Offline Spike Sorter (version 3.3, Plexon Inc., Dallas, TX, USA), see ***Riehle et al. (2018)***. Synchrofacts, i.e., spike-like synchronous events across multiple electrodes at the sampling resolution of the recording system (1/30 ms) (***Torre et al., 2016)***, were then removed. Sorted units were separated into broad- and narrow-spiking SUs representing putative excitatory and inhibitory neurons, respectively. The separation was achieved by thresholding the spike-widths distribution (***Bartho et al., 2004***; ***Kaufman et al., 2010, 2013***; ***Peyrache et al., 2012***; ***Dehghani et al., 2016)*** in the following way. For a given monkey, average waveforms from all SUs recorded in all considered sessions (REST and R2G) were collected. Based on the distribution of spike-widths (time interval between trough and peak of a waveform), thresholds for “broadness” and “narrowness” were chosen such that the values in the middle of the distribution stayed unclassified (Fig. 7). For monkey N, spikes with a width shorter than 0.4 ms were considered to be narrow (ns—narrow-spiking SUs, putative inhibitory neurons), whereas spikes longer than 0.41 ms were considered to be broad (bs—broad-spiking, putative excitatory neurons). For monkey E, spikes narrower than 0.33 ms were considered as ns SUs and spikes broader than 0.34 ms were considered as bs SUs. The difference between monkeys was due to different filter settings during the recordings.

Next, a two step classification was performed. For a given session, the thresholds were applied to the *averaged* SU waveforms (first preliminary classification). Secondly, the *single* waveforms of all SUs were thresholded and a consistency measure *c* was calculated per SU: the percentage of SU single waveforms preliminarily classified as broad. If *c>* 0.5, a SU was classified as bs; if *c<* 0.5,a SU was classified as ns (second preliminary classification). Typically, these two classifications yielded inconsistent results for some single units, e.g., a SU with majority of spikes slightly narrower than 0.4 ms has been classified (based on its *average* waveform) as bs SU. During an iterative procedure we increased the minimal required consistency until there were no more contradictions in the results of both preliminary classifications. SUs with high enough consistency were then declared classified as putative excitatory or inhibitory. SUs whose mean waveform’s widths fell between two thresholds or whose consistency was too low were declared unclassified.

Only SUs with signal-to-noise ratio (SNR, see ***Hatsopoulos et al. (2004)***) of at least 2.5 and a minimal average firing rate of 1 Hz were considered for the analysis to ensure enough and clean data for valid statistics.

### Behavioral Segmentation

Based on video recordings, each REST session was segmented according to monkey’s behavior. Three states were defined with single-second precision as follows: resting state (RS)—no movements and eyes open; sleepy resting state (RSS)—no movements and eyes (half-)closed; and spontaneous movements (M)—accounting for movements of the whole body and limbs (Fig. 1A). If a movement or a RSS interval began during a particular second, this whole second was classified as M or RSS, respectively. Eye movements and minor head movements were allowed during RS. All other types of behaviour (e.g., strong isolated head movements) and periods for which it was not possible to clearly classify the monkey’s behavior (e.g., due to a lack of visibility) were considered as unclassified and excluded from analyses. To increase the reliability of classification, behavioral scoring for each session was performed by two independent observers, and the results were merged later.

Behavior classification in R2G recordings was based on the events registered throughout the experiment, see Sec. Materials and Methods: Experimental paradigm and recordings. Two periods within a trial (see Fig. 1B) were considered: preparatory period (PP), defined as 500 ms after the CUE-OFF (first half of the preparatory delay, no movements), and task-related movement (TM)— 500 ms after SR-ON, including grasping. Due to differences in performance speed of each monkey, this period was defined as: [SR-ON, SR+500 ms] for monkey E, and [SR-150 ms, SR+350 ms] for monkey N. All successful trials were used.

Since the amount of data strongly differs between behavioral states in REST, we used data slices of equal length, mostly 3 s slices, to have comparable statistics. The choice of slice length represents a compromise between different factors: a) the slice length cannot exceed the typical duration of each behavior (shortest for movement), b) the slice length should be as long as possible for reliable estimation of covariances within each slice, c) to average across slices, we need as many slices as possible. Following these arguments, each behavioral segment was cut into as many continuous slices as possible. For example, if a REST segment was 7 s long, it was separated into two slices of 3 s and the remaining 1 s was not considered for the analysis. In the R2G data, the slice length for the 2 behavioral states was 0.5 s by definition, see above. When directly comparing REST and R2G data, we either considered 0.5 s slices for the REST data (comparison of firing statistics) or we concatenated six 0.5 s slices of the R2G data to 3 s slices (analysis of covariances and balance).

#### LFP Spectra

The spectral density of the LFP (sampling frequency of 1000 Hz) in different behavioral states in REST and R2G data was estimated with Welch’s method provided by Elephant (https://python-elephant.org). We considered 3 s slices for the REST and 0.5 s slices for the R2G data. The spectra shown in Fig. 1 were obtained by averaging over single spectra from state-specific slices of all respective recordings. We used a Hanning window of 1 s and an overlap of 50% for the REST data while the R2G spectra were estimated with a Hanning window of 0.3 s with an overlap of 50%. Additionally, an artifact in session N1—high-amplitude synchronous peak on all recording channels—was removed: it was replaced by the average of the remaining signal.

### Data analysis

To characterize and compare different behavioral states, we employed a set of analysis tools. We quantified the correlation between neuronal firing and behaviour and characterized firing properties of SUs, as well as the coordinated firing of pairs of SUs. We also calculated the dimensionality of spiking and the balance between time-resolved putative excitatory and inhibitory population counts.

Pre-processing and data analyses were performed in Python, version 2.7, with the Elephant pack-age (https://python-elephant.org). Since our distributions were typically non-Gaussian, significance of differences between them was assessed via Kruskal-Wallis tests for multivariate comparisons (KW, non-parametric alternative to a one-way ANOVA), with significance level *α* = 0.001. Multiple comparisons were corrected for with a Bonferroni-Holm correction.

For visualizations of distributions obtained for different behavioral states, we used notched boxplots. The line in the center of each box represents the median, box’s area represents the interquartile range, and the whiskers indicate minimum and maximum of the distribution (outliers excluded).

#### Behavioral correlation

For each REST session, we defined a state vector based on the behavioral segmentation, see Sec.Materials and Methods: Behavioral Segmentation. Each entry represented 1 s of the recording and was set to −1 for RS, to −2 for RSS and to 1 for M. To assess the relation between SU activity and monkey’s behavior, the FR (in 1 s bins, no overlap) of each SU was correlated (Spearman rank correlation) with the modified state vector of a given session: entries for RSS were zeroed. Only pairs of entries in which the modified state vector was different from zero were considered. This procedure resulted in a value which we called behavioral correlation: BC ∈ [-1,1], and the corresponding *p* value (indicating statistical significance if *p<* 0.001, with correction) for each SU, see Fig. 2B. Positive BC indicated that the FR increased during movements or decreased during rest, and vice versa. We investigated the distributions of BC values separated between ns and bs SUs, see Fig. 2C.

For a substantiation of these results, we additionally performed Kruskal-Wallis tests on all three behavioral states defined in REST (M, RS, and RSS), separately for each SU, to check for significant changes in the SU firing rates. Note that this method does not provide any quantification of amplitude of the correlation similar to BC.

For both tests described above, we calculated the percentage of SUs that changed their FR significantly (after correction) with changes in the behavioral states, see Tab. 2. This procedure was performed for all SUs and separately for ns and bs neurons.

Next, we again applied Kruskal-Wallis tests for pairwise comparisons between three behavioral states per SU, asking for a significant in- and decreases in firing. This analysis was performed on 3 s long data slices. Tab. 3 lists the percentages of all SUs which either significantly increased or decreased their FR in one state with respect to another.

#### Neuronal firing in REST and R2G states

To compare the SU firing properties in behavioral states from different experiments, we used 0.5 s-long slices of both REST and R2G recordings. In REST, the single seconds at the transitions from one state to another were excluded. For each time slice of each SU, we estimated the average firing rate FR and the local coefficient of variation CV2 (***Ponce-Alvarez et al., 2010***; ***Voges and Perrinet, 2010***; ***Riehle et al., 2018)***, and per SU across slices the Fano factor FF (***Nawrot et al., 2008***; ***Nawrot, 2010***; ***Riehle et al., 2018)***. For the REST recordings, we also calculated the commonly used coefficient of variation CV (***Shinomoto et al., 2003***; ***Ponce-Alvarez et al., 2010***; ***Voges and Perrinet, 2010)***, shown in the additional Figure in Appendix 1 Sec. Transient activities.

CV and CV2 are based on the inter-spike-interval distribution of a SU (***Shinomoto et al., 2003***; ***Ponce-Alvarez et al., 2010***; ***Voges and Perrinet, 2010***; ***Riehle et al., 2018)***. They characterize the (ir-)regularity in neuronal firing. A value close to zero indicates regular spiking, a value of one indicates Poissonian spiking, and a value above one even more irregular firing. The CV2 corrects for transient firing rate changes which yield inappropriately high CV values (***Ponce-Alvarez et al., 2010***; ***Voges and Perrinet, 2010)***. The FF describes the variability in SU spike counts across trials (R2G) or time slices (REST) (***Nawrot et al., 2008***; ***Nawrot, 2010***; ***Riehle et al., 2018)***.

We compared the FR and CV2 values obtained for each SU within each slice of RSS, RS, M, TM and PP states in two different ways, see Fig. 3. On the one hand, we averaged over time slices/trials to represent the variability with respect to SUs. On the other hand, we averaged the results obtained for each data slice/trial over SUs in order to analyse the variability of our measures in time. The significance of the differences between the behavioral states was assessed with a Kruskal-Wallis test including a Bonferroni-Holm correction, both when comparing all 5 states and in pairwise comparisons.

#### Covariances and dimensionality

To measure the joint variability in rate modulation, we calculated the pairwise spike-count co-variances (COV, ***Cohen and Kohn (2011)***; ***Dahmen et al. (2019)***). REST data were cut and R2G data concatenated into 3 s slices (state-resolved) and binned into 100 ms intervals. The bin size of 100 ms was a compromise between obtaining enough bins to calculate covariance values (given a slice length of 3 s), considering enough spikes for reliable estimation of covariance, and using a time scale appropriate for the examination of rate modulations. For R2G data this procedure implied that data from 6 consecutive trials contributed to a single COV value.

The COV between spike trains *i* and *j* was defined as:

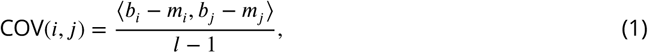

with and —binned spike trains, and being their mean values, *l* the number of bins, and ⟨*x, y*⟩ the scalar product of vectors *x* and *y*. Thus, for each 3 s slice of a particular state we obtained a COV matrix 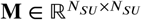 with *N*_*SU*_—number of SUs.

Based on the COV matrices, we calculated the participation ratio PR to characterize the dimensionality of activity in different behavioral states, see ***Mazzucato and La Camera (2016)***; ***Gao et al. (2017)***. Eigenvalue decomposition of COV matrix **M** yields *N*_*SU*_ eigenvalues *λ* with corresponding eigenvectors *v*, such that **M***v*_*i*_ = *λ*_*i*_*v*_*i*_. The eigenvalues were used to calculate the participation ratio of the neuronal dynamics:

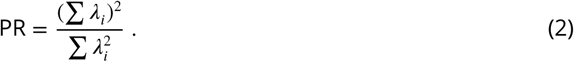

The PR thus quantifies how many eigenvectors are necessary to explain a significant part of variance in dynamics described by **M**, see Fig. 8A.

The PR is low if most of the variability is captured by the first few eigenvectors. A large PR indicates that many eigenvectors are necessary to capture the dynamics—a sign of high complexity. In order to test the robustness of our results, we performed our analysis with different bin sizes. The result is shown in (Fig. 8B). Here, all bin sizes revealed the same PR-dependent ordering of behaviors. This suggests that our results are robust to the choice of bin size.

The value of the PR depends on the number of SUs present in the analysis: It can take values 1≤ PR ≤*N*_SU_. In order to make the PRs comparable across recording sessions, we normalize with the number of SUs measured in the respective experiment, PR^′^ = PR /*N*_SU_, thus giving rise to values in a range [0, 1]. The box plots in Fig. 4 visualize the distributions of normalized PRs obtained in all non-overlapping 3 s time slices from all recording sessions of a given type (REST or R2G) per monkey. The time-resolved visualization of PR^′^ in panel B of the same Figure was calculated in a sliding-window fashion with 3 s slices and 2 s overlap. Each data point in this plot is located at the center of the respective window.

#### Balance

The multiscale balance between putative excitatory (bs SUs) and inhibitory (ns SUs) population firing was examined similarly to the procedure proposed in ***Dehghani et al. (2016)***. We considered timescales from 1 ms to 10 s. For a given timescale, pooled spikes from bs and ns units were binned and z-scored separately, resulting in bs and ns population activities (whole recording, no separation into behavioral states). Then, the putative inhibitory population activity was subtracted from the putative excitatory activity (see Fig. 5A). If this difference was close to zero, i.e., if pooled ns and bs spike counts were nearly identical, the network activity was called balanced. If this was the case for multiple time scales (i.e., bin sizes), it was called multiscale balance.

Since we observed some deviations from balance for bin sizes larger than 30 ms, we quantified these deviations in a state-resolved manner. For each REST and R2G session of a given monkey, we binned the 3 s time slices (concatenated from six consecutive trials of 0.5 s for R2G data) into 100 ms bins. Next, we applied two methods to quantify the balance between population activities. Firstly, the same as for the multiscale balance, we z-scored the population activities, using the respective mean and standard deviation of the whole recording (not state-specific). Then, we calculated, separately for each state, the difference between the z-scored bs and ns population activity of each 100 ms bin in each time slice: A negative value indicated a domination of ns activity while a positive value meant that the bs activity was higher. Fig. 5C shows the corresponding state-resolved histograms.

Secondly, we calculated the Spearman rank correlation between raw bs and ns population activities for each time slice: The higher the correlation *ρ*(bs, ns), the more strict the instantaneous balancing between the ns and bs populations (cf. (***Renart et al., 2010***; ***Tetzlaff et al., 2012)***). The state-resolved results are presented in box plots (Fig. 6A).

To investigate the relationship between balance and dimensionality, we calculated the Spearman rank correlation between *ρ*(bs, ns) and PR^′^ for each monkey, pooled over all REST and R2G sessions, respectively (Fig. 6C, D and Tab. 6).

## Acknowledgments

This project has received funding from the Deutsche Forschungsgemeinschaft Grant GR 1753/4-2 & DE 2175/2-1 Priority Program (SPP 1665), the Helmholtz Association through the Helmholtz Portfolio Theme Supercomputing and Modeling for the Human Brain (SMHB), and from the European Union’s Horizon 2020 Framework Programme for Research and Innovation under Specific Grant Agreement No. 720270 & 785907 (Human Brain Project SGA1 & SGA2). We thank Tom Tetzlaff and Alper Yegenoglu for fruitful discussions.

## Appendix 1 Transient activities

To analyze the instantaneous balance, we correlate putative excitatory and inhibitory population activities in different behavioral states (sleepy rest RSS, rest RS, and movements M). We find a significantly reduced correlation (i.e., balance) during RS compared to M for monkey E, and during RS and M compared to RSS in monkey N, significantly only between M and RSS (cf. 6A and B). We also observe numerous transient increases in the population spike counts (Fig. 5B). Such simultaneous peaks contribute to higher correlation values between the two neuronal populations. The prevalence of this deviations differs between behavioral states. Fig. 1 and Tab. 1 below show that the distributions of population activities during M (monkey N, ns population) or both RSS and M (monkey E, both populations) are characterized by higher standard deviations than expected from higher mean values. RSS of monkey N shows lower means and slightly higher standard deviations than RS in ns population, pointing to the same conclusion. Both relations serve as footprints of an increased number of narrow peaks in population spiking during non-resting states.

Given the transient peaks in the population spike counts during M and RSS, we suspect the following relationship between balance and transients in the population activity: Whenever one of the population activities transiently increases, the other one is forced to do the same due to the recurrent coupling between putative excitatory and inhibitory neurons, yielding higher correlation value and thus more balance.

**Appendix 1 Figure 1.**
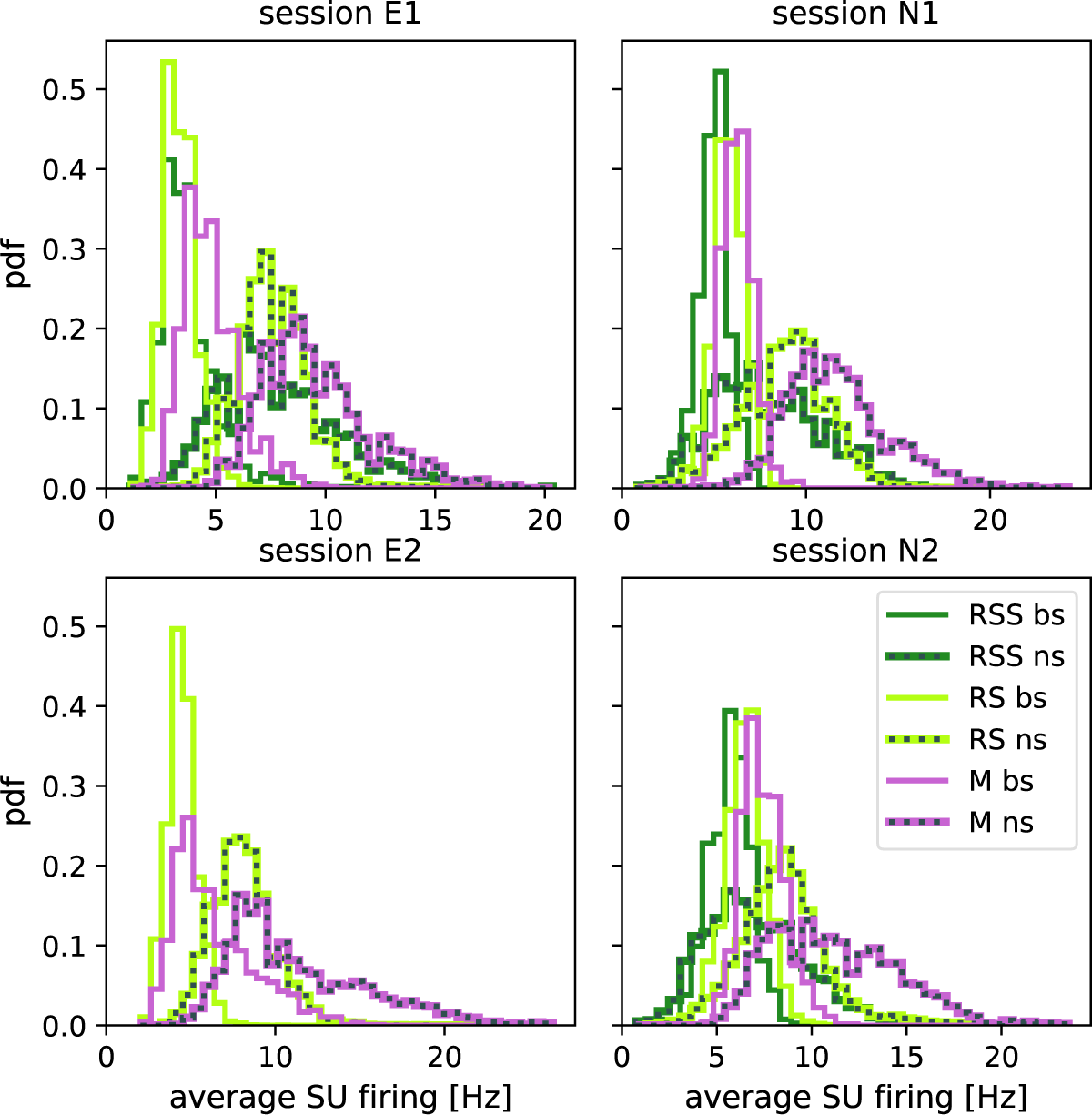
Population spiking in REST recordings. Distributions of the population spiking activities calculated in 100 ms bins, separately for each REST session (left column—monkey E, right—monkey N, colors indicate behavioral states, solid lines—bs, dashed lines—ns). Notice different ranges of the x axes.

**Appendix 1 Table 1.**
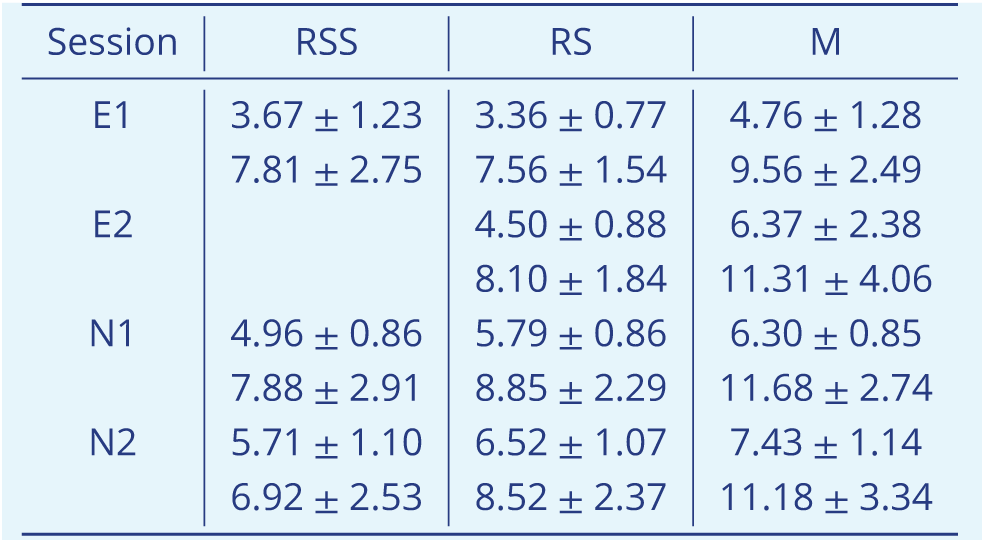
Mean values and standard deviations of distributions of population spiking activities visualized in Fig. 1 above. First line per session: bs, second line: ns population.

Complementary to the above discussed population activity, we now look at the firing on the level of SUs (considering 3 s time slices), shown in the top row of Fig. 2. We expect higher firing rates (mean and standard deviation) for states with more transient activities (M&RSS) which show an increased balance in the population activities. For monkey E, the reduced balance in RS compared to M (cf. Fig. 6A) coincides with a significant average firing rate reduction in RS compared to M. However, for monkey N, the reduced balance in M compared to RSS (cf. Fig. 6B) coincides with a significant firing rate increase in M compared to RSS. Since SU firing rate cannot be directly related to instantaneous balance, we examine another feature of firing, namely its regularity. The bottom row of Fig. 2 compares the results obtained for two different regularity measures, CV and CV2: only CV2 accounts for transient firing rate changes which typically yield erroneously high CV values (***Ponce-Alvarez et al., 2010***; ***Voges and Perrinet, 2010)***. Thus, a significant difference between CV and CV2 suggests the presence of such transient firing rate changes. Indeed, we observe significantly higher CV during RSS and M. Obviously, this is no proof but only an indication for transient changes in the firing rates on the level of SU firing during movements and sleepy rest.

**Appendix 1 Figure 2.**
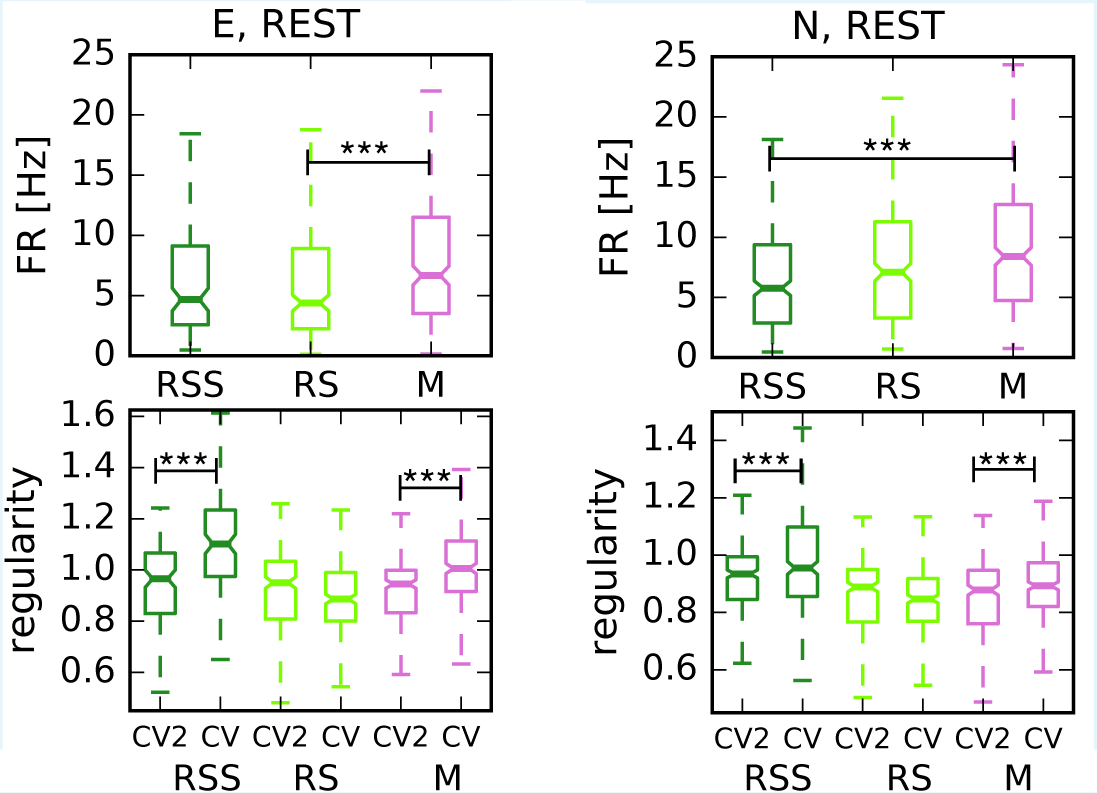
Transient activities in REST recordings. Box plots of average firing rates (top row) and two different regularity measures (bottom row) of the three states defined for REST recordings (RSS in dark green, RS in light green, and M in magenta) of monkey E (left) and monkey N (right). CV2 and CV characterize the (ir-)regularity in spiking, but only CV2 accounts for transient changes in the firing rates which typically yield misleadingly high CV values. In both monkeys, CV yields significantly higher values than CV2 during M and RSS states but not during RS.

## Appendix 2 Spatial activity distribution

The participation ratio quantifies the dimensionality in the network activity space. One could ask how this relates to the distribution of neuronal activity in physical space. In analogy to large-scale resting state studies which find widely distributed networks of brain areas that are particularly active during rest (***Biswal et al., 1995***; ***Raichle, 2009***; ***Deco et al., 2011)***, we estimated the spatial spread of active SUs in the different behavioral states of the REST recordings. To this end, we calculated the average spatial distance from each active SU to the center of mass of the spiking activity during sleepy rest (RSS), rest (RS), and movements (M). An *active* SU emitted at least one spike during the respective 3 s slice; the center of mass is given by the average coordinates of all active SUs. We thus characterized the mean spatial spread of the activity around the center of mass in each behavioral state.

Figure 1 (below) shows the results obtained for our four REST sessions. The distinct scales on the y-axis are a result of the different implantations of the Utah arrays in the two monkeys (number and placement of active versus inactive electrodes), see ***Riehle et al. (2018)***. The differences in the spatial confinement of active SUs were small but consistent across sessions and monkeys. For monkey E, we find that the activity during RS exhibits a higher spatial spread than during M, even if the difference is only weakly significant (*p<* 0.01 in session E1 and *p <* 0.05 in session E2). In monkey N, only the second session shows a significant difference, namely a larger spatial spread in RS compared to M (*p <* 0.01). In summary, we show a tendency of the SU activity during rest to be distributed over a larger spatial region than during movement, which may relate to higher dimensionality quantified by PR.

**Appendix 2 Figure 1.**
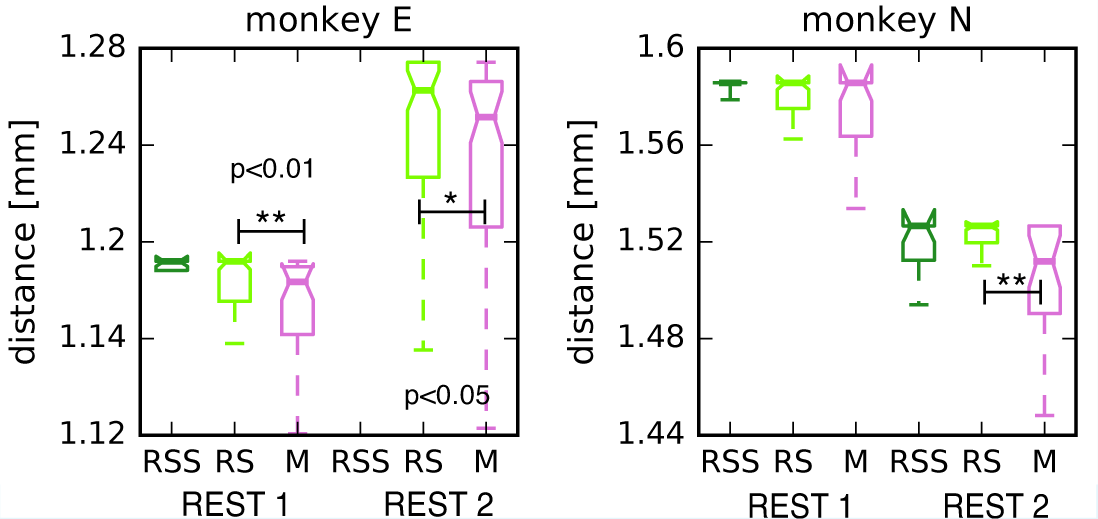
Spatial arrangement of active SUs on the Utah array. Box plots of the radial spatial distance to the center of mass for the two REST recording sessions of monkey E (left panel) and N (right panel).

## Footnotes

1 Our results are less significant than those presented in ***Riehle et al. (2018)***; we analyze only a subset of the R2G data and use partially different methods.

2 For example, PP is defined as 500 ms after CUO-OFF when the monkey is forbidden to move, and constraining the analyzed data to only successful trials ensures that the monkey was focused on the upcoming cue to perform the task.

